# Inducible mismatch repair streamlines forward genetic approaches to target identification of cytotoxic small molecules

**DOI:** 10.1101/2023.02.21.529401

**Authors:** Thu P Nguyen, Min Fang, Jiwoong Kim, Baiyun Wang, Elisa Lin, Vishal Khivansara, Neha Barrows, Giomar Rivera-Cancel, Maria Goralski, Christopher L Cervantes, Shanhai Xie, Johann M Peterson, Juan Manuel Povedano, Monika I Antczak, Bruce A Posner, David G McFadden, Joseph M Ready, Jef K De Brabander, Deepak Nijhawan

**Affiliations:** Department of Biochemistry, University of Texas Southwestern Medical Center, Dallas, TX 75390, USA; Quantitative Biomedical Research Center, Department of Clinical Sciences, University of Texas Southwestern Medical Center, Dallas, TX 75390, USA; Department of Internal Medicine, Program in Molecular Medicine and Division of Hematology/Oncology, University of Texas Southwestern Medical Center, Dallas, TX 75390, USA; Department of Internal Medicine, Program in Molecular Medicine and Division of Endocrinology, University of Texas Southwestern Medical Center, Dallas, TX 75390, USA; Simmons Cancer Center, University of Texas Southwestern Medical Center, Dallas, TX 75390, USA

## Abstract

Orphan cytotoxins are small molecules for which the mechanism of action (MoA) is either unknown or ambiguous. Unveiling the mechanism of these compounds may lead to useful tools for biological investigation and in some cases, new therapeutic leads. In select cases, the DNA mismatch repair-deficient colorectal cancer cell line, HCT116, has been used as a tool in forward genetic screens to identify compound-resistant mutations, which have ultimately led to target identification. To expand the utility of this approach, we engineered cancer cell lines with inducible mismatch repair deficits, thus providing temporal control over mutagenesis. By screening for compound resistance phenotypes in cells with low or high rates of mutagenesis, we increased both the specificity and sensitivity of identifying resistance mutations. Using this inducible mutagenesis system, we implicate targets for multiple orphan cytotoxins, including a natural product and compounds emerging from a high-throughput screen, thus providing a robust tool for future MoA studies.

## Introduction

Understanding the mechanism of action for anti-cancer small molecules is a critical milestone in drug development and remains a significant challenge. Small molecules bind to many proteins depending on the concentration used, therefore, it is challenging to precisely pinpoint which interaction is driving a specific phenotype. Considered “gold standard” criteria, a target protein is one for which a point mutation is identified that does not alter protein function but confers resistance to a compound both in a cell-free biochemical assay as well as in a cellular context.^1^ Such a mutation might reduce the affinity to the compound while not critically reducing the activity of the protein.

Even when there is detailed structural information that illustrates how a compound and protein interact, it is difficult to nominate a mutation that meets these criteria. An alternative strategy is to subject a target protein to random mutagenesis within its native cellular context and select for a specific phenotype, such as resistance to compound treatment. Numerous technologies have been used to mutagenize a gene of interest and identify resistance mutations, including CRISPR-Cas9 nuclease and base-editor tiling screens, or expression constructs mutagenized through error-prone PCR or propagation in mutator strains of *E. coli*.^2–5^ These approaches are limited in scope and inherently biased as they rely on prior knowledge of a hypothetical target.

Forward genetic screens represent an unbiased approach and have been successfully used to identify resistance mutations under circumstances when a compound’s target is unknown. A requirement of forward genetics is the ability to mutagenize a population before selecting for a phenotype of interest. The colorectal cancer cell line, HCT116, which has an elevated mutation rate due to defects in DNA mismatch repair, has been used as a forward genetic tool to identify resistance mutations for anti-cancer compounds.^6^ The emergence of resistant cells in HCT116 following compound selection is both more frequent and more likely to be the result of mutation when compared to mismatch repair-proficient cells. Moreover, HCT116 cells are diploid, which leads to an enrichment for heterozygous mutations like those in the binding site of inhibitors. Previously, our group utilized HCT116 to discover mutations that cause either indisulam or CD437 resistance in cells and block compound activity *in vitro*, thus fulfilling the gold standard target identification criteria for these compounds.^7,8^ In both cases, whole exome sequencing of several independent resistant clones revealed recurrent binding site mutations, suggesting an exceedingly high mutation rate in the population and tolerance for amino acid substitutions at key residues. Although these examples illustrate the power of HCT116 forward genetics for target identification, the number of successful cases remains limited, suggesting technical challenges in obtaining clonal resistance and identifying causal mutations.

The enhanced mutagenesis found in HCT116 represents a double-edged sword – the high mutation rate, which may be essential for generating rare variants, also leads to numerous background mutations that make it challenging to identify causal variants. A recent evaluation of the mutation rate in HCT116 cells using next generation sequencing technology reported a base pair mutation rate of 46 x 10^-7^ (including both base pair and frameshift), which is 11,740-fold higher than the overall mutation rate during normal hematopoiesis of 3.9 x 10^-10^.^9,10^ Based on these estimates, the expansion of 3 million cells over three doublings will yield 24 million cells with 3.4 x 10^9^ exome mutations, more than 100-fold coverage of the human exome. These predictions are consistent with our observation that rare variants are recurrently found in HCT116 cells and explain the high number of background mutations. Unfortunately, for many orphan compounds, genetic heterogeneity in resistance, such as mutations in multiple residues across a large protein or mutations in different genes, might make it difficult to distinguish causal mutations from background under these conditions. Thus, the current methodology may not successfully identify these mutations, and may in part explain the limited success of HCT116 cells in revealing compound-resistant mutations.

Here, we describe the development of isogenic cancer cell lines, including HCT116, in which we temporally control mutagenesis by regulating the levels of DNA mismatch repair proteins. We demonstrate that this isogenic cell platform enables more efficient and sensitive identification of compound resistant mutations by i) assigning the compound resistance phenotype to increased rates of mutagenesis; ii) precisely defining ideal selection concentrations; and iii) improving causal mutation calling by reducing background mutation noise and expanding the types of mutations that can be readily assigned to compound resistance. We successfully used this platform to unveil resistance mutations for seven different compounds, thus highlighting the utility of this approach in target identification efforts.

## Results

### Engineering inducible mutagenesis for compound resistance mutation discovery

The first step in developing a system with fewer background mutations is to engineer the ability to temporally control the mutation rate of the cell. Since the high mutation rate in HCT116 is the result of loss of MLH1 protein, we set out to engineer an HCT116 clone in which we could temporally regulate the levels of MLH1.

First, we tested whether ectopic expression of MLH1 is sufficient to reduce the clonal resistance rate of HCT116 cells. Expression of MLH1 in HCT116 led to increased levels of its heterodimeric binding partner, PMS2, suggesting that ectopically expressed MLH1 is functional (Figure 1A).^11^ We next assessed whether MLH1 re-expression suppressed the emergence of compound resistant clones by exposing HCT116^WT^ and HCT116^MLH1^ cells to a lethal dose of MLN4924, a small molecule inhibitor of NEDD8-activating enzyme (NAE) that acts by binding to its catalytic subunit UBA3. ^12^ Following selection of one million cells with MLN4924, 14 clones emerged from the HCT116^WT^ cells, but none were identified in the HCT116^MLH1^ cells (Figure 1B). Five out of six sequenced HCT116^WT^ clones harbored a UBA3^A171T^ mutation, consistent with a previous report showing that UBA3^A171T^ mutation renders cells resistant to MLN4924 toxicity (Supplement figure 1A and B). ^13^ Taken together, we concluded that exogenous expression of MLH1 was sufficient to suppress mutagenesis and the emergence of compound resistant clones in HCT116.

**Figure 1.**
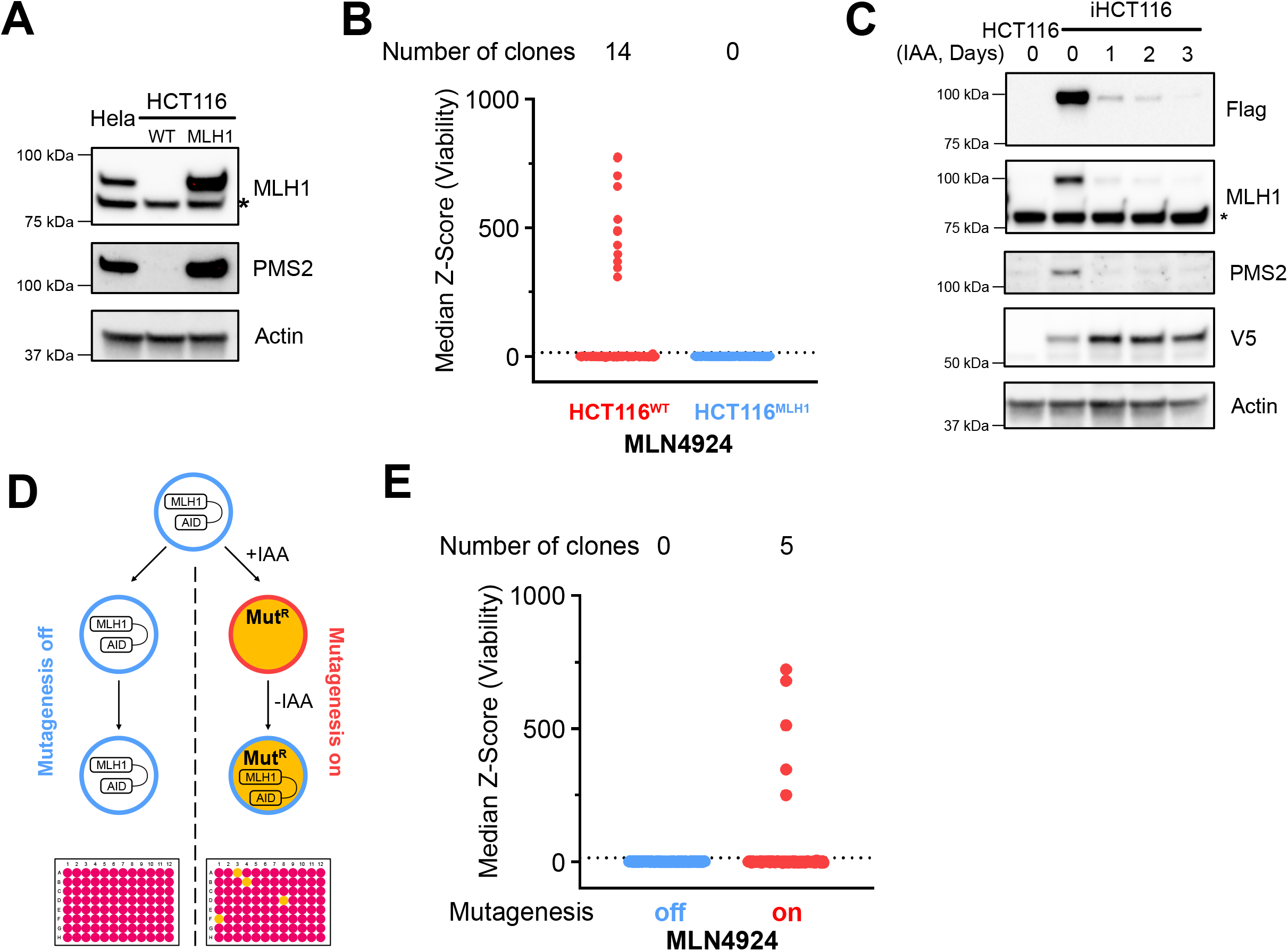
Engineering inducible forward genetics system for compound resistance mutation discovery. (A) Western blot of lysates derived from Hela, HCT116, and HCT116-MLH1. (B, E) Cell viability. Outliers (clones) defined as median z-Score >15 (dashed line). (C) Western blot of lysates derived from HCT116 and iHCT116 treated with 500 μM indole-3-acetic acid (IAA). (D) Inducible forward genetics system screening schematic. Asterisk (*) denotes non-specific band.

Having demonstrated that MLH1 expression is sufficient to reduce resistance rates, we expressed a form of MLH1 that could be temporally regulated by the addition of auxin, a non-toxic small molecule. In plants, auxin (indole-3 acetic acid; hereafter referred to as IAA) acts as a molecular glue that recruits auxin-inducible degron (AID) domains to the E3 ubiquitin ligase TIR1, leading to the ubiquitylation and proteasomal degradation of AID domain-containing proteins.^14^ IAA can be used to regulate the levels of mammalian proteins by ectopically expressing TIR1 and tagging the intended target protein with an AID domain.^15^ We ectopically expressed MLH1 fused to a C-terminal AID domain and Flag epitope tag, as well as TIR1 (with a V5 epitope tag), in HCT116 cells and isolated a single clone hereafter referred to as iHCT116. Exposure of iHCT116 to IAA led to the rapid and durable degradation of MLH1 (Figure 1C). We next tested whether induced MLH1 depletion and DNA mismatch repair deficiency was sufficient to promote the emergence of compound resistant clones. iHCT116 cells were cultured in the absence (Mut-off) or presence (Mut-on) of IAA for two weeks to accumulate mutations. Following mutagenesis, we reasoned that mutations were fixed in the population, thus, we withdrew IAA prior to compound selection to restore MLH1 expression and reduce the cellular mutation rate (Figure 1D). Following exposure of 500,000 cells from each population to a lethal dose of MLN4924, the Mut-on population yielded five resistant clones while the Mut-off population had none (Figure 1E). Six clones from a separate selection were isolated and further analyzed for their sensitivity to MLN4924. Compared to the parental iHCT116, all six clones were resistant to MLN4924, with an increase in IC_50_ of at least 35-fold (Supplement figure 1C), and five out of six clones harbored UBA3^A171T^ mutations (Supplement figure 1D). These experiments demonstrate that inducible degradation of ectopic MLH1 in HCT116 can yield clones that harbor compound-resistant mutations in the target.

### Target deconvolution for PVHD303, a tubulin binder with a putative alternative target

We first applied the iHCT116 platform to two previously reported scaffolds that have compelling *in vitro* evidence supporting a target, but ambiguity regarding the cellular mechanism. In such cases, compound-resistant mutations have the potential to provide critical biological insight. PVHD121 and its more potent derivative, PVHD303, (Figure 2A) are quinazoline derivatives that compete for colchicine binding to tubulin and inhibit tubulin polymerization *in vitro*. ^16,17^ The cellular effects of these compounds, however, are thought to be distinct from colchicine, which has raised the possibility that there is an alternative cellular target.^17,18^ Cells treated with relatively high concentrations of PVHD121 or PVHD303 have abnormal spindles and no microtubules, consistent with inhibition of tubulin polymerization. However, at concentrations close to the anti-proliferative IC_50_ of the respective compounds, cells exhibit a more specific defect in centrosome-specific micronucleation. ^17,18^ These differences could be explained by either a different binding mode or an alternative target, and the identification of compound resistant mutations using iHCT116 might help differentiate between these two possibilities.

**Figure 2.**
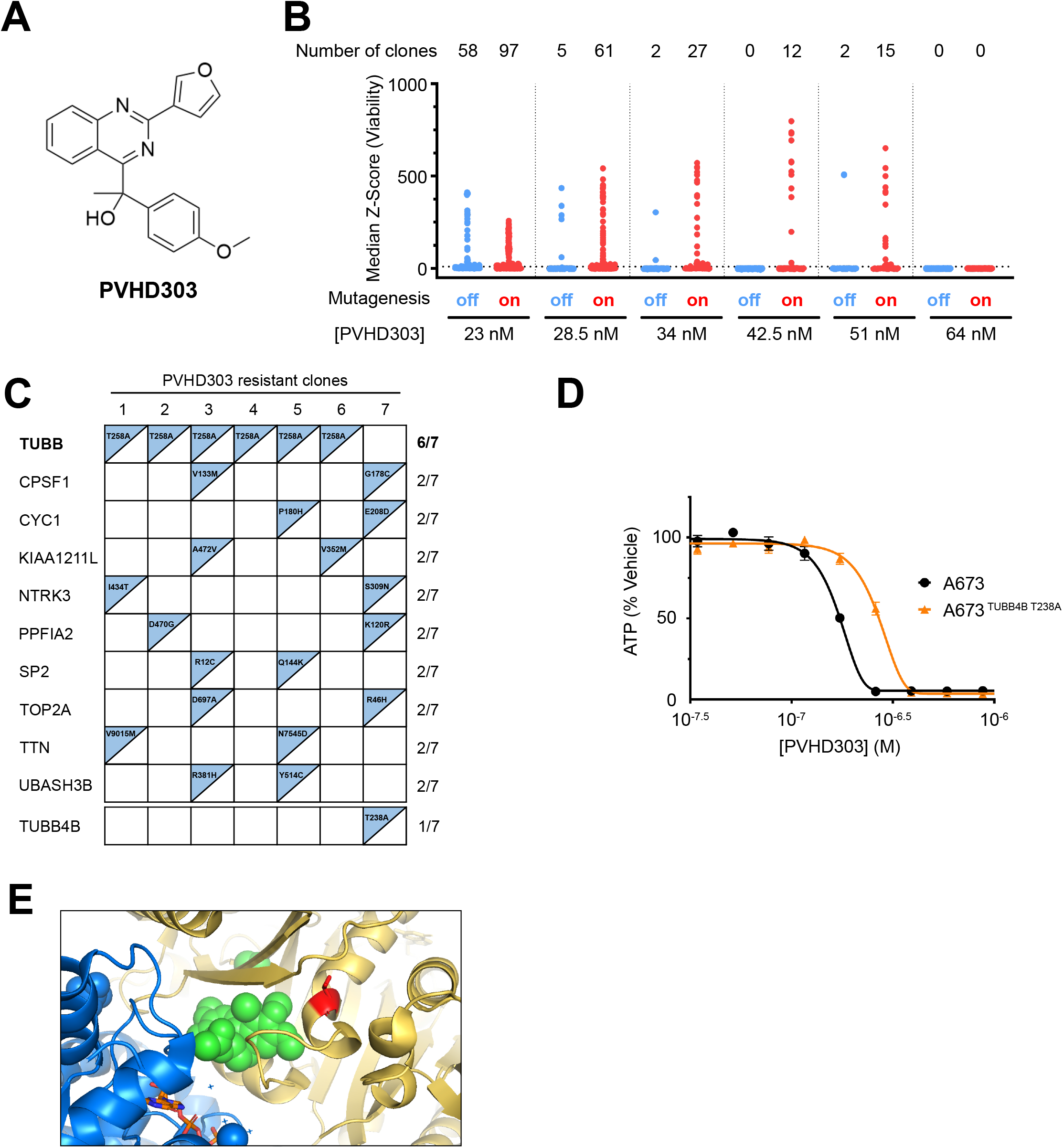
Mutation in tubulin renders cells resistant to PVHD303. (A) PVHD303. (B) Cell viability. Outliers (clones) defined as median z-Score >10 (dashed line) (C) Whole-exome sequencing (WES) analysis results for PVHD303 resistant clones. (D) Viability assay of A673^WT^ and A673^TUBB4B T238A^ cells treated with PVHD303 in dose-response format. Points and error bars represent mean +/- SEM of two technical replicates. Error bars are not shown if SEM is smaller than the point. (E) T258 residue (red) is mapped to tubulin co-crystal structure with Colchicine (PDB: 6XER). Blue is α-tubulin, yellow is β-tubulin, GTP (orange), and colchicine (green).

Considering that PVHD303 may have two anti-proliferative targets, we reasoned that the potency of each target would differ, even if by a small degree. Concentrations of compound that engage both targets are not expected to yield clonal resistance because of the low likelihood of two compound-resistant mutations in a single clone. Therefore, we selected for clonal resistance at several concentrations ranging from 23 nM to 64 nM. The number of surviving clones was higher in the Mut-on conditions at doses between 23 nM and 51 nM. When selected with 23 nM PVDH303, the ratio of Mut-on/Mut-off clones was 97:58, and this ratio increased to 15:2 when selected with 51 nM PVDH303. These results suggest that at lower concentrations, there are background surviving clones, which may not be the result of mutation. There were also fewer total clones at the 51 nM concentration, which we estimate reflects reduced genetic heterogeneity. It is worth noting that at slightly higher concentrations (64 nM), no clones were present (Figure 2B).

Following the analysis of barcode sequences, seven independent clones were expanded for further investigation. All clones were resistant to PVHD303, with a 1.76- to 2.76-fold increase in IC_50_ (Supplement figure 2A). Interestingly, all seven clones were equally sensitive to colchicine, suggesting that the resistance mechanism was specific to PVHD303 (Supplement figure 2B). We performed whole-exome sequencing (WES) on these 7 clones and the parental iHCT116, and for each clone we identified any mutation present in the clone but not the parental population. Of note, we excluded 104 genes that were recurrently mutated in clones selected against independent compounds (Supplement table 1). Some of these genes, like *MUC2* and *MUC6*, harbored repetitive sequences that may have contributed to a high mutation rate. ^19^ After removing these genes, we listed any gene that was recurrently mutated in two or more clones raised from each compound.

One gene – *TUBB* – which encodes an isoform of β-tubulin, was mutated in six out of seven clones. All six of these clones harbored TUBB^T258A^ mutations, and the remaining clone harbored a homologous mutation (TUBB4B^T238A^) in a different β-tubulin isoform (Figure 2C). We reasoned that these mutations, albeit in different isoforms, were functionally equivalent, and set out to determine whether TUBB4B^T238A^ mutations cause resistance to PVDH303. To demonstrate causality, we introduced a biallelic TUBB4B^T238A^ mutation into A673 cancer cells using CRISPR-Cas9 (Supplement Figure 2C). A673 cells expressing TUBB4B^T238A^ were resistant to PVHD303 in comparison to wild-type A673 cells (Figure 2D). TUBB^T258A^ (and the homologous TUBB4B^T238A^) localized to the colchicine binding pocket (Figure 2E), consistent with a previous report showing PVHD303 competed with colchicine in a binding assay. Cells harboring TUBB4B^T238A^, however, were not resistant to colchicine (Sup Figure 2D), suggesting a different binding mode. Taken together, these results demonstrated that the anti-cancer activity of PVHD303 was rescued by tubulin mutations and suggested that the centrosome specific micronucleation was either the consequence of how this molecule binds to tubulin or is the result of binding another target which does not cause cell death at the most potent (lowest) concentrations.

### Identifying resistance mutations to guide target campaigns for PARP-1 binders

Poly [ADP-ribose] polymerases, or PARPs, are a family of 17 enzymes that catalyze the transfer of ADP-ribose from NAD^+^ to a substrate. The clinical success of PARP-1 inhibitors such as olaparib, niraparib, and rucaparib in the treatment of cancer has garnered interest in the potential clinical utility of targeting other PARP enzymes. The biological function of PARPs has been implicated in mitosis, and therefore, one hypothesis is that inhibition of PARPs may lead to defects in mitosis such as multipolar spindles (MPS). ^20,21^ With this rationale, AstraZeneca screened a library of phthalazinone derivatives (presumably synthesized as part of a PARP-1 lead optimization program) to identify molecules that induce MPS, reasoning that one or more of these molecules would inhibit other PARPs. Indeed, they identified AZ9482, a phthalazinone derivative that induced MPS and caused cell death (Figure 3A).^22^ Even though AZ9482 inhibited PARP-1 *in vitro*, many other equally potent PARP-1 inhibitors did not induce MPS or cause cell death. Therefore, the induction of MPS and cell death did not correlate with PARP-1 inhibition *in vitro*, raising the possibility of an alternative cellular target. Although PARP-6 was implicated as the functional target based on a correlation between *in vitro* inhibition and cellular activity, subsequent chemical proteomic experiments did not support this target.^23,24^ To date, there have been no genetic approaches used to identify the target of AZ9482.

**Figure 3.**
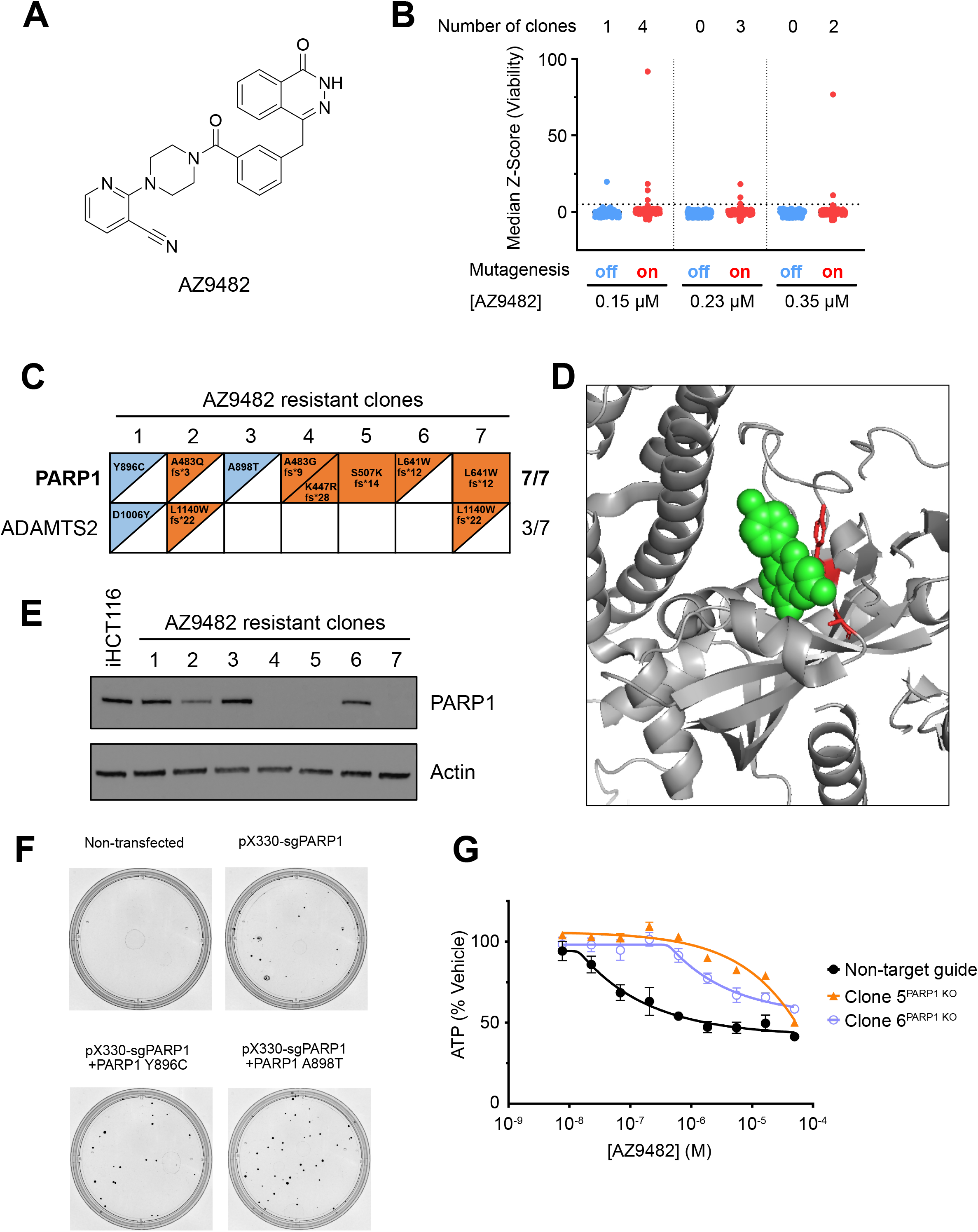
Mutations in PARP1 render cells resistant to AZ9482. (A) AZ9482. (B) Cell viability. Outliers (clones) defined as median z-Score >5 (dashed line). (C) WES analysis results for AZ9482 resistant clones. (D) Y896 and A898 (red) residues are mapped on PARP-1 (gray) co-crystal structure with Talazoparib (green) (PDB:4UND). (E) Western blot of lysates derived from iHCT116 and AZ9482 resistant clones. (F) Crystal violet stain of transfected HCT116 cells following 350 nM AZ9482 treatment. (G) Viability assay of HCT116 harboring non-target guide or PARP1 KO clones.

To identify compound resistance mutations, we treated mutagenized and unmutagenized populations of iHCT116 with lethal doses of AZ9482. Mut-on populations consistently yielded more clones than Mut-off populations when cells were treated with 0.15, 0.23, or 0.35 μM AZ9482 (Figure 3B). Seven independent clones were expanded, tested for AZ9482 resistance, and analyzed by WES (Supplement figure 3A). In our analysis, which included all types of mutations, only 2 genes were mutated in more than 2 clones – *PARP-1* (mutated in all 7 clones) and *ADAMTS2* (3 clones) (Figure 3C). Clone 1^AZ9482^ and Clone 3^AZ9482^, harbored the missense mutations PARP-1^Y896C^ and PARP-1^A898T^, respectively. These heterozygous mutations resided in the catalytic domain near the binding site of the PARP-1 inhibitor talazoparib (Figure 3D). ^25^ The remaining 5 clones contained insertion-deletion mutations in one or both alleles of *PARP-1*, resulting in the introduction of frameshifts and premature stop codons (Figure 3C). While the missense mutations were consistent with mutations that affected compound binding, the identification of nonsense mutations suggested that reduced PARP-1 protein levels might cause resistance. Indeed, we observed no PARP-1 protein in AZ9482 Clones 4, 5, and 7, and reduced PARP-1 levels in AZ9482 Clones 2 and 6 (Figure 3E). To test whether these genetic events caused AZ9482 resistance, we employed CRISPR-Cas9 editing to either install the PARP-1^Y896C^ or PARP-1^A898T^ mutations, or silence PARP-1. In the case of the former, cells were transfected with a non-targeting single guide RNA (sgRNA) or an sgRNA targeting *PARP-1*, in the absence or presence of a single-stranded DNA repair template that encoded the Y896C or A898T mutations. Following selection with 350 nM AZ9482, clonal resistance was substantially higher in cells treated with both the repair template and *PARP-1* sgRNA (Figure 3F). Similarly, two clones isolated from a population of cells infected with *Cas9-sgPARP-1*, which had significantly lower levels of PARP-1, were resistant to AZ9482 treatment (Figure 3G, Supplement figure 3B). Although these results did not resolve the mechanism underlying MPS induction, our data demonstrated that the anti-proliferative activity of AZ9482 was due to its effect on PARP-1.

### Identifying resistance mutations to better understand the mechanism of natural products as exemplified by Destruxins

Genetic approaches to target identification, unlike chemical proteomics, do not necessarily require any additional chemical derivatives or chemical optimization. This advantage is particularly relevant for natural products, which often have complex structures that are not readily amenable to modification for chemical proteomics. Therefore, we applied the iHCT116 platform to elucidate the cellular mechanism of the Destruxins, a class of depsipeptides secreted by the fungus *Metarhizium anisopliae* with multiple different biological activities. ^26^ Two Destruxins – B and E (hereafter abbreviated as DB and DE) – have been shown to inhibit the proliferation of colorectal cancer cells and impair xenograft tumor growth. ^27,28^ Beyond these anti-proliferative effects, DB and DE have been reported to inhibit bone resorption by inducing morphological changes in osteoclasts, suppress Hepatitis B virus replication in cell culture models, and inhibit T-lymphocyte mediated cytotoxicity. ^29–32^ One proposed cellular target for the Destruxins is the vacuolar-type ATPases (V-ATPases). V-ATPases are large, multi-subunit complexes composed of a V_0_ domain that traverses the membrane and a V_1_ domain. The V_1_ domain catalyzes ATP hydrolysis to promote proton flux through a channel formed by the V_0_ domain; these concerted actions lead to the acidification of cellular organelles like the lysosome.^33^ Inhibitors of the V-ATPase include bafilomycin, which inhibits proton transport and reduces lysosomal acidification by binding to a V_0_ domain subunit.^34^ Previous work demonstrated that Destruxin B inhibits proton transport by the V-ATPase *in vitro*, and that DB and DE inhibit lysosomal acidification in cells. ^35,36^ The current hypothesis is that Destruxins target the V-ATPase, but several questions remain including which, if any, of the bioactivities is due to V-ATPase inhibition, and whether the mechanism of V-ATPase inhibition is distinct from other inhibitors.

DB and DE are similar in structure, except that the propyl side chain in DB is replaced by an epoxy group in DE (Figure 4A). We used DE for clonal selection because anti-proliferative effects were achieved at substantially lower concentrations than DB (Supplement figure 4A). Mut-on and Mut-off iHCT116 populations were subjected to increasing concentrations of DE ranging from 77 nM to 116 nM. We observed more clones in the Mut-on population following selection with 77 nM and 96.5 nM DE, suggesting that mutations were driving resistance. Following exposure to 116 nM DE, no clones were isolated from either population, suggesting that the mutations isolated at lower selection doses had low penetrance (Figure 4B). We evaluated six independent clones for sensitivity to either DE or DB, and, compared to parental iHCT116, all six clones were resistant to DE, with an increase in the IC_50_ of 2.5- to 3.7-fold (Supplement figure 4B). Four clones (Clone 2^DE^, Clones 4-6^DE^) showed cross-resistance with DB, but surprisingly, Clone 1^DE^ and Clone 3^DE^ were more sensitive to DB (Supplement figure 4C). Our WES analysis pipeline identified mutations in *ATP6V1B2*, a subunit of the V_1_ domain, in five out of six resistant clones. Clone 1^DE^ and clone 3^DE^, which were both more sensitive to DB, harbored an ATP6V1B2^H384Y^ substitution. Clone 2^DE^, Clone 4^DE^, and Clone 6^DE^ displayed R381K, A194T, and D359N mutations, respectively (Figure 4C, supplement 4D). Although these mutations are separated within the ATP6V1B2 primary structure, all these mutations map in close physical distance within the V_0_V_1_-ATPase cryo-EM structure, suggesting that this region of the protein is important for compound activity (Figure 4D). To test whether these mutations caused resistance, we used CRISPR-Cas9 editing to install the ATP6V1B2^D359N^ mutation into compound-naïve HCT116 cells. Cells were transfected with either a non-targeting sgRNA or an sgRNA targeting *ATP6V1B2*, in the absence or presence of a single-stranded DNA repair template that encodes for the D359N mutation. Following selection with 96 nM DE, clonal resistance was substantially higher in cells treated with both the repair template and *ATP6V1B2* sgRNA (Supplement figure 4E). We expanded the surviving cells from the ATP6V1B2^D359N^ knock-in selection (hereafter referred to as HCT116^ATP6V1B2 D359N^) and assessed sensitivity to DB and DE. Compared to wild-type HCT116, HCT116^ATP6V1B2 D359N^ cells were resistant to both DB and DE (wild-type: DB IC_50_ = 0.780 μM, DE IC50 = 0.0354 μM; ATP6V1B2^D359N^: DB and DE IC50 > 10 μM) (Figure 4E). Wild-type HCT116 and HCT116^ATP6V1B2 D359N^ cells were comparably sensitive to taxol, an unrelated toxin, and bafilomycin A1, suggesting that ATP6V1B2^D359N^ specifically blocked the Destruxin mechanism (Figure 4E, Supplement figure 4G).

**Figure 4.**
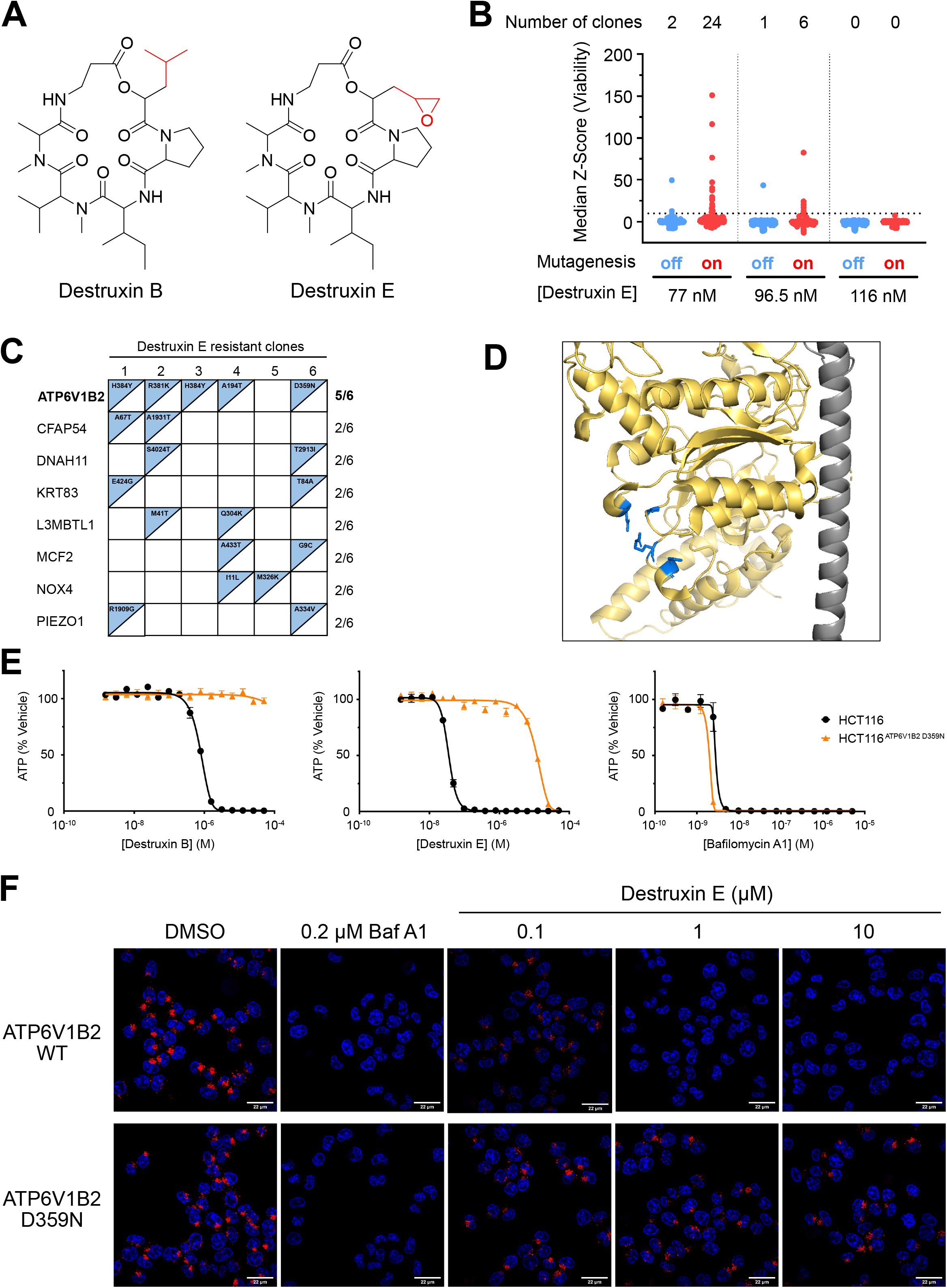
Mutations in ATP6V1B2 renders cells resistant to destruxin B and E. (A) Destruxin B and destruxin E. (B) Cell viability. Outliers (clones) defined as median z-Score >10 (dashed line). (C) WES analysis results for Destruxin E resistant clones. (D) Compound resistant mutations are mapped to v- ATPase crystal structure (PDB: 6WM2). Gray is ATP6V1E1, yellow is ATP6V1B2. The mutated residues are highlighted in blue. (E) Viability assay of HCT116^WT^ and HCT116^ATP6V1B2 D359N^ treated with destruxin B, destruxin E, or Bafilomycin A1 in dose-response. Points and error bars represent mean +/- SEM of two technical replicates. Error bars are not shown if SEM is smaller than the point. (F) Confocal images of HCT116^WT^ and HCT116^ATP6V1B2 D359N^ treated with bafilomycin A1 or destruxin E. Nuclei are blue. Lysotracker is red. Bar-scale represent 22 μm.

These findings supported the hypothesis that the anti-cancer activity of the Destruxins is through V-ATPase inhibition. To further test this hypothesis, we assayed whether DB and DE inhibit V-ATPase activity in cells. Wild-type and ATP6V1B2^D359N^ HCT116 cells were treated with increasing concentrations of DB and DE and stained with LysoTracker, a pH-sensitive dye used to visualize acidic cellular compartments. Destruxin treatment caused a dose-dependent decrease in LysoTracker signal in wild-type HCT116 cells, consistent with V-ATPase inhibition, while HCT116^ATP6V1B2 D359N^ cells retained LysoTracker signal following exposure to all doses of DB and DE (Figure 4F and Supplement Figure 4H). Combined, these results demonstrate that DB and DE inhibit the V-ATPase and that this inhibitory mechanism is mediated by targeting ATP6V1B2.

It was notable that the degree of resistance in the ATP6V1B2^D359N^ knock-in population was substantially higher than the original isolated clones. One explanation for this difference is that the original clones possessed a wild-type allele whereas CRISPR-Cas9 cutting may have generated indels that silenced the wild-type allele in the knock-in population. Consistent with this hypothesis, deep sequencing of the region spanning the ATP6V1B2^D359N^ mutation revealed the D359N substitution in approximately 72% of alleles, with a collection of indels/frameshifts comprising the remaining fraction of alleles (Supplement figure 4F). These results suggest that most cells were compound heterozygotes for ATP6V1B2^D359N^ and a silencing frameshift, which resulted in mono-allelic expression of ATP6V1B2^D359N^. Taken together, these results suggest that ATP6V1B2^D359N^ was entirely resistant to the compound but was haploinsufficient in the presence of a wild-type ATP6V1B2 allele. In addition, these experiments provide an example of how iHCT116 can be used to unveil the anti-cancer mechanism for a natural product that is reported to have many different cellular activities.

### Target deconvolution for compounds emerging from high throughput screens

After demonstrating the iHCT116 platform could be used to deconvolute the cellular target for compounds with a proposed MoA, we next sought to apply this strategy for target identification of compounds with no reported mechanism. We screened a chemical library of 99,599 synthetic compounds at a single concentration (2.5 μM) for anti-proliferative activity to HCT116 cells. In a confirmation screen, 862 putative HCT116 toxins were re-analyzed for anti-proliferative activity at multiple concentrations. Based on the results of the confirmation screen, as well as a review for chemical tractability and commercial availability, 89 compounds were selected for further investigation using the iHCT116 system. 36 of these compounds were either cytostatic or inadequately toxic to perform selections, and the remaining 53 compounds were screened for IAA dependent clonal resistance at multiple concentrations. 13 compounds yielded at least two-fold more clones in Mut-on conditions when compared to Mut-off conditions (Supplement figure 5A).

Of these 13 compounds, MM201, MM202, and MM204 are three chemically distinct compounds that were screened for mutation-dependent resistance independently and found to have the same target (Figure 5A). Following selection with lethal doses of each compound, we observed substantially more clones in the MLH1-depleted condition, suggesting that resistance was mutagenesis-driven (Figure 5B). We expanded resistant clones from each selection, assayed sensitivity to MM201, MM202, and MM204, and observed that, compared to parental iHCT116, the tested clones were resistant to their respective selection compounds (Supplement figure 5B). Following WES of six MM201 resistant clones, three MM202 resistant clones, and six MM204 resistant clones, we obtained a short list of putative targets for these compounds which included nicotinamide phosphoribosyltransferase (NAMPT), which was mutated in 6/6 MM201 clones, 3/3 MM202 clones, and 5/6 MM204 clones (Figure 5C).

**Figure 5.**
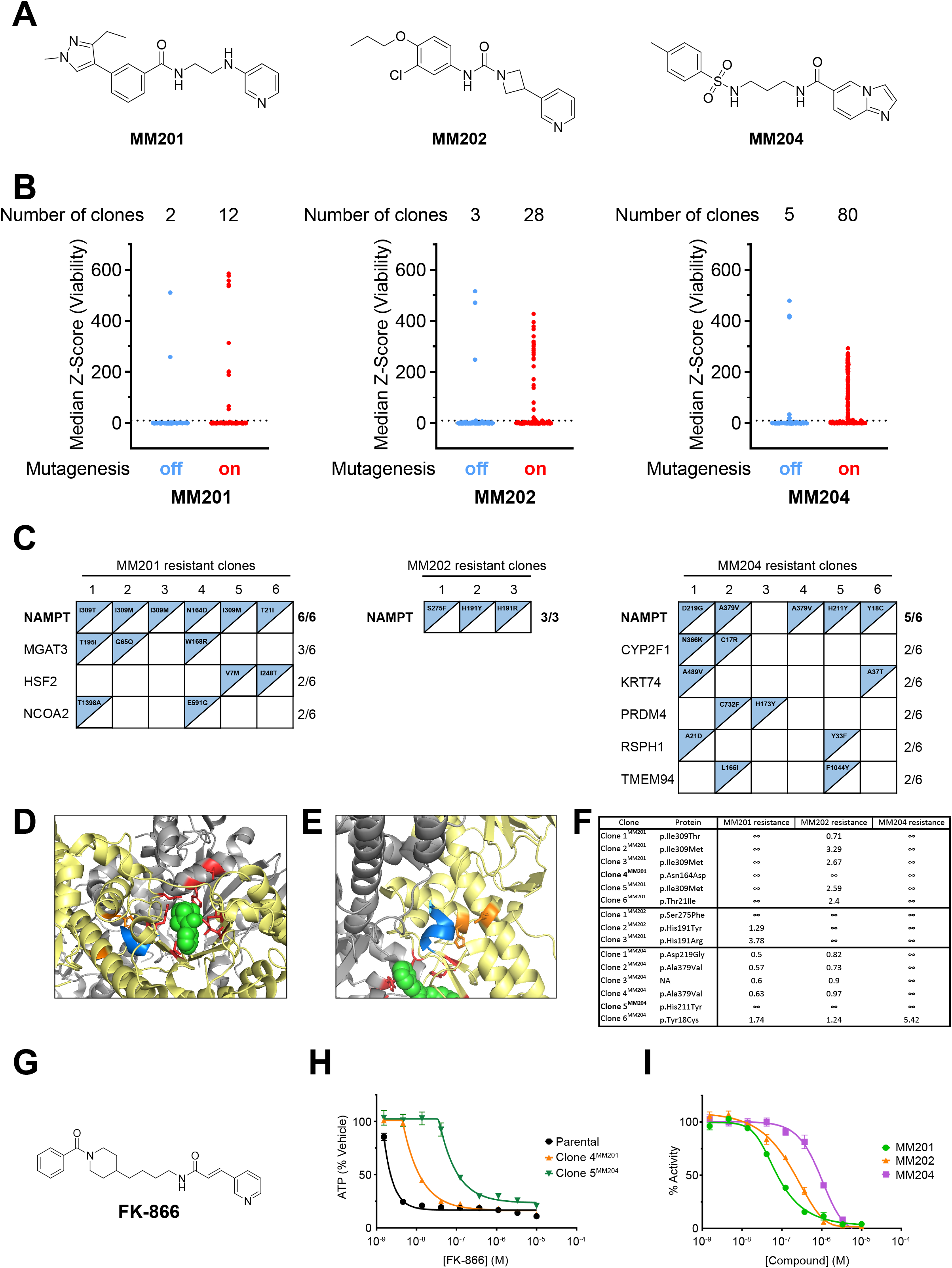
Forward genetics reveals multiple chemical scaffolds targeting Nicotinamide phosphoribosyltransferase (NAMPT). (A) MM201, MM202, and MM204. (B) Cell viability. Outliers (clones) defined as median z-Score >10 (dashed line). (C) WES analysis results for MM201, MM202, and MM204 resistant clones. (D, E) Compound resistant mutations (red or orange) are mapped to NAMPT co-crystal structure with FK-866 (green) (PDB: 2GVJ). Gray and yellow are 2 chains forming the homodimer. G_383_GGLLQ_388_ helical motif is blue. (F) Summary of all clone sensitivity against MM201, 202, and 204. Values reported are fold-resistance in comparison to parental iHCT116. (G) FK-866. (H) Viability assay of HCT-116, clone 4^MM201^, and clone 5^MM204^ treated with FK-866 in dose-response. Points and error bars represent mean +/- SEM of two biological replicates. Error bars are not shown if SEM is smaller than the point. (I) *in vitro* NAMPT activity assay in the presence of MM201, MM202, or MM204 in dose-response. Points and error bars represent mean +/- SEM of three biological replicates. Error bars are not shown if SEM is smaller than the point.

The distribution of NAMPT missense mutations associated with each compound was as follows – MM201: T21I (1 clone), N164D (1 clone), I309M (3 clones), I309T (1 clone); MM202: H191R (1 clone), H191Y (1 clone), S275F (1 clone); MM204: Y18C (1 clone), H211Y (1 clone), D219G (1 clone), A379V (2 clones). Although resistant clones raised against the three compounds shared no common mutations, most of the mutations clustered in the NAMPT pocket that binds a well-characterized inhibitor, FK-866 (Figure 5D, 5G). ^37^ Examination of the NAMPT crystal structure revealed that the N164D and H211Y mutations were adjacent to the G_383_GGLLQ_388_ helical motif, which is a common phosphate-binding motif in various proteins (Figure 5E). Prior work demonstrated that two missense mutations – NAMPT^S165F^ and NAMPT^S165Y^ – destabilized this α-helix, which impairs phosphoribosyl pyrophosphate (PRPP) binding.^38^ Since many NAMPT inhibitors need to be phosphoribosylated to achieve their cellular potency, the loss of this short α-helix, though tolerable for cell growth, might cause compound resistance.^38^ Given the proximity of N164 and H211 to the G_383_GGLLQ_388_ α-helix, we speculated that these mutations worked in a similar manner. Indeed, Clone 4^MM201^ and Clone 5^MM204^, which harbor N164D and H211Y mutations, respectively, showed cross-resistance to FK-866, MM201, MM202, and MM204 (Figure 5F, H; Supplement figure 5C).

The NAMPT reaction utilizes PRPP and ATP to convert nicotinamide to nicotinamide mononucleotide (NMN). In addition to NAMPT’s contribution to NMN synthesis, cells can generate NMN from nicotinamide riboside (NR) through a reaction catalyzed by nicotinamide riboside kinase 1 (NMRK1) (Supplement Figure 5D). If a compound’s cytotoxic activity is through NAMPT inhibition and impaired synthesis of NMN, then restoring NMN levels through supplementation with exogenous NR should rescue cell viability following compound treatment. Consistent with this hypothesis, we observed that FK-866 toxicity was rescued by supplementation with increasing doses of NR. Additionally, NR rescued the toxicity of MM201, MM202, and MM204, suggesting that the anti-proliferative effect of these compounds is through NAMPT inhibition (Supplement Figure 5E). Finally, we utilized an *in vitro* biochemical assay to determine that MM201, MM202, and MM204 directly inhibit NAMPT with IC_50_’s of 0.0743, 0.177, and 0.952 μM, respectively (Figure 5I). Combined, these results provide genetic and biochemical evidence that MM201, MM202, and MM204 are diverse chemical scaffolds that inhibit NAMPT *in vitro* and in cells. Moreover, this example demonstrates how iHCT116 can be used to efficiently guide target identification studies for compounds with unique chemical composition that directly emerge from a high-throughput screen.

SW394703 is another small molecule identified as an HCT116 toxin (Figure 6A). While SW394703 was toxic to HCT116 cells (IC_50_ = 1.21 μM), its enantiomer, SW394704, was inactive, even at concentrations as high as 50 μM (Figure 6A, Supplement figure 6A). Selections with SW394703 yielded multiple resistant clones that were raised in a mutation-dependent manner (Figure 6B). Six independent clones were assayed for compound sensitivity, and we found that all six clones were resistant to SW394703 (Supplement Figure 6B). Through WES, we identified five genes that were mutated in more than one clone, and one gene – *DDB1* – that was mutated in four independent clones (Figure 6C). Given that the mutations in *DDB1* were predicted to be loss-of-function (nonsense mutations or frameshift mutations causing premature stop codons), we evaluated DDB1 protein levels and found that clones with *DDB1* mutations had lower levels of DDB1 (Figure 6D). These results suggested that reduced DDB1 protein levels confer resistance to SW394703.

**Figure 6.**
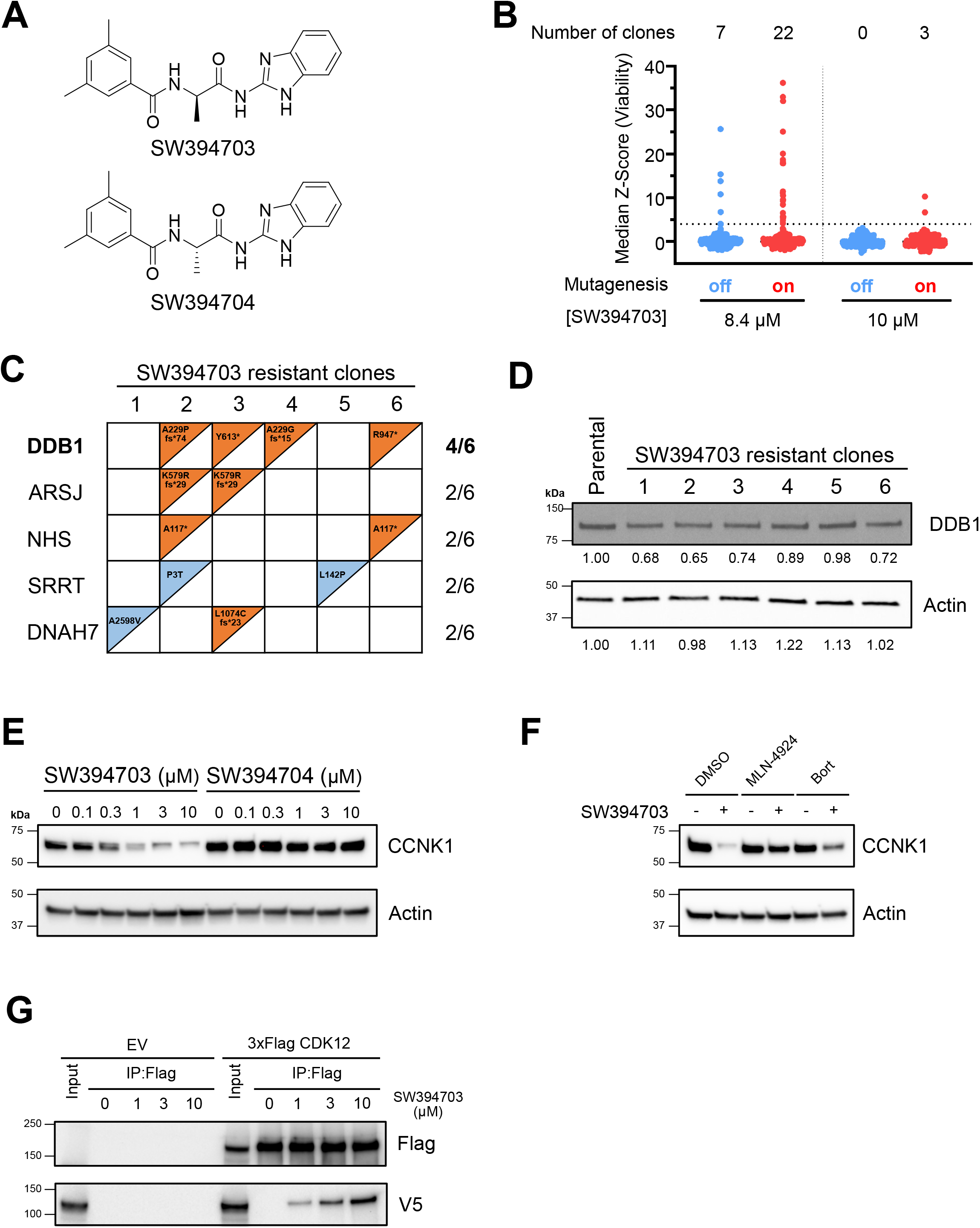
Loss-of-function mutations in DDB1 render cells resistant to SW394703. (A) SW394703 and SW394704. (B) Cell viability. Outliers (clones) defined as median z-Score >4 (dashed line). (C) WES analysis results for SW394703 resistant clones. (D) Western blot of lysates derived from iHCT116 and SW394703 resistant clones. (E, F) Western blot of lysates derived from iHCT116 treated with SW394703 or SW39704 in the absence or presence of MLN4924 or Bortezomib pretreatment. (G) Immunoprecipitation and western blot of lysates derived from vector or 3xFlag-CDK12 expressing HCT116, in the presence or absence of SW394703.

DDB1 is an essential adapter protein in cullin ring ligase 4 (CRL4) ubiquitin ligase complexes and acts by bridging substrate receptors to the cullin 4 scaffold protein. ^39,40^ A genetic relationship between *DDB1* and SW394703 resistance led us to hypothesize that SW394703 might function as a molecular glue degrader. Molecular glue degraders are a class of small molecules that act by recruiting neo-substrates to ubiquitin ligases leading to neo-substrate ubiquitylation and proteasomal degradation. ^41^ In some cases, it has been shown that the expression level of specific ubiquitin ligase components directly correlates with the degree of neo-substrate degradation. ^8^ As part of a strategy to identify anti-cancer small molecules that work as glue degraders, Ebert and colleagues found that the anti-cancer activity of CR8, a cyclin dependent kinase inhibitor, correlated with the expression of DDB1. ^42^ Subsequent studies demonstrated that CR8 recruits CDK12 to DDB1, which leads to the ubiquitylation and proteasomal degradation of the CDK12 heterodimeric partner, cyclin K. In two independent studies, at least four more structurally diverse small molecules were shown to have the same activity, raising the possibility that CDK12/cyclin K recruitment to DDB1 is promiscuous. ^43,44^

Based on our observations, and the results of prior studies, we hypothesized that SW394703 may act through a mechanism similar to CR8. Treatment with SW394703, but not its stereoisomer, led to a reduction in cyclin K protein levels at concentrations consistent with the anti-proliferative activity of SW394703 (Figure 6E). Additionally, SW394703-induced cyclin K degradation was blocked by co-administration of MLN4924, which inhibits neddylation and cullin-mediated ubiquitylation, or the proteasome inhibitor bortezomib (Figure 6F). To further understand the mechanism of SW394703-mediated cyclin K degradation, we assessed whether SW394703 induces a complex between DDB1 and CDK12. Lysates were collected from cells engineered to ectopically express V5-tagged DDB1 and Flag-tagged CDK12, incubated with increasing doses of SW394703, and CDK12-containing complexes isolated by immunopurification. We observed that SW394703 promoted the dose-dependent formation of a CDK12-DDB1 complex that was not observed with the inactive enantiomer, SW394704 (Figure 6G and Supplement figure 6C). Finally, SW394703 resistant clones were cross-resistant to CR8 (Supplement figure 6D). These genetic and biochemical data suggest that SW394703 is a new chemical scaffold that recruits CDK12 and cyclin K to DDB1, leading to cyclin K degradation in a similar mechanism to the previously reported molecular glue degraders CR8, HQ461, and dCeMM2/3/4. ^42–44^

### Development of an inducible forward genetic system in a cell line with intact mismatch repair

Initially, forward genetic studies with cancer cells were restricted to HCT116 or other cells that have intrinsic mismatch repair deficiencies. We previously demonstrated that silencing of MSH2, an essential component of DNA mismatch repair, promotes increased mutagenesis and enables forward genetic screening in cancer cell lines that are otherwise mismatch repair proficient. ^45^ Having demonstrated the utility of inducible mismatch repair defects in HCT116, we sought to establish an analogous system in S462 cells, a human malignant peripheral nerve sheath tumor (MPNST) cell line with no known defect in mismatch repair. We first evaluated whether silencing MSH2 with CRISPR-Cas9 was sufficient to increase the clonal resistance rate to MLN4924 in S462 cells. Populations of wild-type and MSH2-null S462 cells were exposed to lethal doses of MLN4924, and the number of surviving clones was determined. Selections in MSH2-null S462 cells yielded four resistant clones per two million cells while no resistant clones were recovered from the wild-type S462 population (Supplement figure 7A and B).

To generate an inducible forward genetics system in S462 (hereafter referred to as iS462), we used CRISPR-Cas9 editing to introduce the sequence encoding an AID domain at the 3’ end of the endogenous *MSH2* locus. Subsequently, TIR1 was ectopically expressed in a clonal *MSH2^AID/AID^* cell line. IAA treatment resulted in the rapid and durable depletion of MSH2 in iS462 cells, which led to the destabilization of its obligate heterodimeric partner, MSH6 (Figure 7A). To assess whether induced degradation of endogenous MSH2 would increase the rate of clonal resistance, iS462 cells were cultured in the presence or absence of IAA and then subjected to selection with a lethal dose of MLN4924. The MSH2-depleted population gave rise to four MLN4924-resistant clones while no clones were recovered from the control population (Figure 7B). Clones isolated from the MSH2-depleted population were tested for sensitivity to MLN4924 and all four clones displayed resistance to MLN4924, with IC_50_ increases of at least 109-fold compared to the parental iS462 cell line. Additionally, these clones all harbored a UBA3^A171T^ mutation (Supplement figure 7C and D). Taken together, these findings demonstrate that mismatch repair-proficient cell lines can be adapted for forward genetics through introduction of inducible mismatch repair deficiencies.

**Figure 7.**
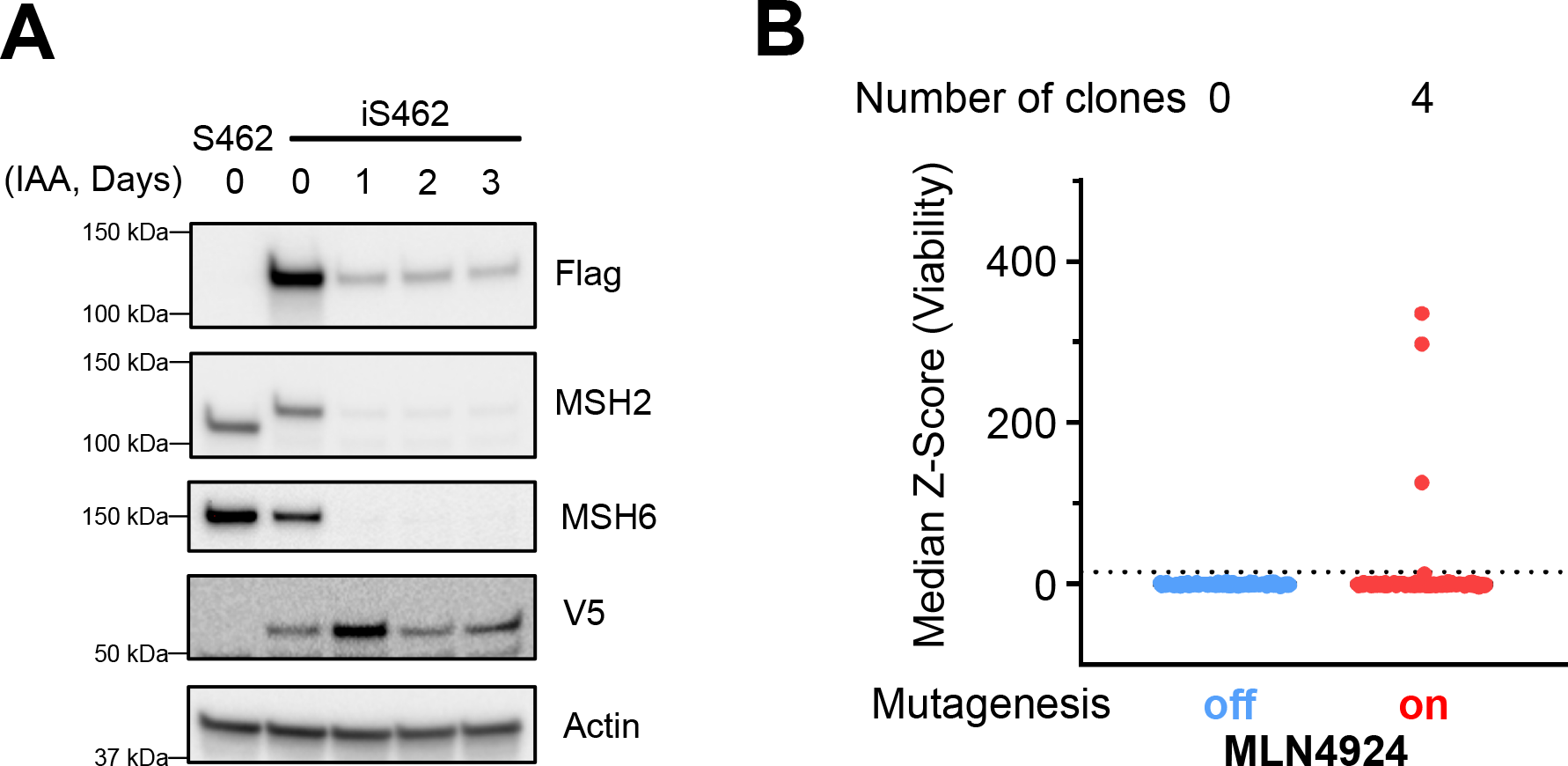
Engineering inducible mutagenesis system in mismatch repair proficient cell line. (A) Western blot of lysates derived from S462 and iS462 treated with 500 μM IAA. (B) Cell viability. Outliers (clones) defined as median z-Score >15 (dashed line).

## Discussion

Unambiguous target identification is required for the interpretation of biological experiments that use small molecule tool compounds and provides critical guidance for the development of small molecule therapeutics. The importance of rigorous target validation in cells is highlighted by several recent examples where compounds were optimized *in vitro* and advanced to the clinic but were later found to work through alternative cellular targets. ^46,47^ For compounds where optimization is guided by an *in vitro* assay, a correlation between the *in vitro* and *in vivo* structure-activity relationship is often utilized to provide evidence for the cellular target. However, confusion may arise if the *in vitro* assay does not faithfully reproduce the effects of cellular target engagement, or the physicochemical properties of a compound influence activity in either the cell or cell-free assay. To overcome this latter caveat, photochemical probes containing alkynes have been combined with medicinal chemistry to evaluate compound-protein interactions in a cellular context. ^48–51^ Notwithstanding, these strategies require extensive chemical optimization in order to build a correlation between structure, binding, and activity. The need for many analogs can limit the applicability or practicality of this approach, especially in the case of compounds such as natural products, which may be difficult to derivatize.

In comparison to biochemical affinity-based strategies, forward genetic approaches to target identification afford several advantages. First, identifying a mutation that confers resistance in a physiologic and cell-free assay provides gold standard evidence of target identification – a level of confidence that biochemical strategies cannot provide. Second, compounds can be screened for their ability to generate resistant clones without further chemical optimization, so this approach can be used for compounds that emerge directly from high-throughput screens, or complex molecules such as natural products. Third, compound-resistant mutations provide rigorous controls for defining on-target effects in a physiologic context. Finally, in the case of small molecule drugs, compound-resistant mutations may provide insight into potential mechanisms of resistance and guide the development of next-generation inhibitors. Mismatch repair-deficient cells have been used as forward genetic tools to identify compound-resistant mutations that led to target identification; however, the number of successful examples is limited. Here, we sought to improve this approach by developing isogenic systems in which we conditionally and temporally regulate the levels of mismatch repair proteins and, as a result, the mutation rate.

These isogenic, inducible cell platforms provide a significant improvement to existing forward genetic tools by increasing the sensitivity of resistance mutation detection. The increased sensitivity in mutation calling is driven by the ability to compare isolated clones to a closely paired reference genome (the parental, Mut-off population), which both refines the list of potential causal mutations and expands the types of mutations that can be considered. In our prior studies, due to the high background rate, we were forced to restrict our analysis to missense mutations. Using HCT116 cells, we discovered *RBM39* mutations in approximately one-half of indisulam-resistant clones.^8^ Subsequent biochemical experiments revealed that indisulam is a molecular glue degrader that recruits RBM39 to the E3 ubiquitin ligase CRL4-DCAF15 leading to RBM39 ubiquitylation. Only later did we discover that many of the clones with wild-type RBM39 harbored loss-of-function mutations in different components of CRL4-DCAF15. These mutations initially evaded our detection due to the high background rate and were only found after the mechanism had been determined using other approaches. In this study, we identified compound-resistant clones that harbored missense mutations as well as indels or nonsense mutations. Specifically, loss-of-function mutations in *PARP-1* or *DDB1* caused resistance to AZ9482 or SW394703, respectively, highlighting how inducible mutagenesis can be used to readily identify these resistance mechanisms. Thus, the improved mutation detection sensitivity afforded by these inducible mutagenesis systems may expedite future target identification campaigns by providing multiple genetic clues to a small molecule’s MoA.

Our studies demonstrate that another important advantage of this approach is the ability to isolate mutation-dependent resistant clones over a narrow concentration range. We were able to isolate clones with mutation-driven resistance to PVHD303 and Destruxin E (DE), which afforded only 1.76- to 2.76- fold or 2.47- to 3.66-fold resistance in PVDH303 or DE, respectively. There are multiple conditions in which resistance mutations would have low penetrance including: i) the mutation has a modest effect on compound affinity for the target; ii) there are multiple targets with similar potencies and engagement of the less potent target masks resistance from mutations in the more potent target; or iii) the fraction of protein that is mutated leads to modest resistance, a phenomenon known as allelic dilution. For the Destruxins and PVHD303, the latter is the most likely explanation. When cells were engineered to express only the mutant ATP6V1B2, they were completely resistant to Destruxin treatment, demonstrating that a full complement of the mutant protein is required for penetrant resistance. Similarly, there are multiple isoforms for tubulin, the target of PVHD303, and mutation of a single allele may have influenced only a minor fraction of the cellular tubulin pool. One or more of these scenarios is likely to apply to many orphan cytotoxins, thus, inducible mutagenesis systems may be essential for identifying resistant mutations.

The potential utility of these inducible mutagenesis systems is evident in our ability to rapidly deconvolute the cellular mechanism of compounds with otherwise ambiguous cellular MoA. AZ9482 is a compound for which there was established *in vitro* binding activity, but cellular toxicity was attributed to an alternative target. We found that PARP-1 mutations (including active site mutations) rendered resistance to AZ9482. These findings provide support for the hypothesis that the anti-cancer activity of AZ9482 is in fact due to targeting PARP-1 and suggest that the differential cellular phenotype is rather the result of an alternative binding mode. Different PARP inhibitors have allosteric effects that cause varying degrees of PARP-1 trapping on DNA, which leads to a DNA damage response, resulting in cellular toxicity. ^52^ Our prediction is that AZ9482 causes more PARP-1 trapping, a feature that was not captured with the initial *in vitro* studies. To this end, a correlation between *in vitro* assays for PARP-1 trapping and toxicity may provide further support for the MoA of AZ9482. Currently, lead optimization programs in drug discovery campaigns, especially in the pharmaceutical industry, are guided by *in vitro* assays. As a result, there are likely more examples in which cellular phenotypes do not correlate with *in vitro* activity. Therefore, the application of inducible mutagenesis systems to rapidly identify resistant mutations may guide the development of more informative *in vitro* assays or unveil new cellular targets.

Building on the benefits that these inducible mismatch repair-deficient cell lines provide, we demonstrate that these systems can be used to prioritize compounds identified through viability-based high-throughput screens (HTS) for further study. Small molecule screens that score a loss of viability are often referred to as a “down” assays and are prone to high false positive rates from compounds that cause cell death through non-specific mechanisms such as chemical precipitation or detergent effects. To account for this, many screens are designed to include a secondary screen for biological specificity such as selective activity to a particular cancer lineage, cancer genotype, or biological pathway. While these counter-screens are effective at eliminating non-specific hits, they undoubtedly eliminate biologically useful and interesting small molecules in the process. We propose that the iHCT116 platform provides an alternative counter-screening strategy for hits emerging from anti-proliferative HTS efforts. By providing evidence of genetic resistance, compounds that score in the iHCT116 system are enriched for those that act through a specific protein or protein-protein complex.

While iHCT116 can help capture small molecules with potentially interesting cellular MoAs, additional assays are required to quickly de-prioritize compounds acting on promiscuous targets like mitochondrial proteins in the electron transport chain (ETC) and tubulin. ^49,53–55^ Fortunately, most ETC inhibitors can be eliminated by comparing activity in the presence of glucose or galactose, and chemical probes are now available to evaluate whether molecules engage the colchicine binding site of tubulin. ^50^ Our discovery of three new NAMPT inhibitors with distinct structures suggest that NAMPT is also a promiscuous target. While these compounds all contain a pyridine or imidazopyridine cap group reminiscent of other NAMPT inhibitors, there is significant structural variation amongst the linker and tail fragments of these molecules, suggesting it may be difficult to predict NAMPT inhibitors *in silico*. ^56^ Using the iHCT116 system, we identified missense mutations (NAMPT^N164D^ and NAMPT^H211Y^) that confer resistance to several NAMPT inhibitors; these mutations provide additional counter-screening tools to readily identify and de-prioritize NAMPT inhibitors in HTS campaigns.

Once small molecules acting through promiscuous targets are removed, the streamlined identification of mutations by iHCT116 and other inducible mutagenesis platforms allows for the rapid implication of compound targets, regardless of selectivity. In this regard, the discovery of resistant mutations in an uncharacterized or conventionally undruggable protein may provide the rationale for further investigation. As an example, we used iHCT116 to identify resistance mechanisms for SW394703 and discovered mutations in DDB1, a protein that was, until recently considered undruggable. SW394703 recruits a CDK12:cyclin K complex to DDB1, which leads to cyclin K ubiquityl and proteasomal degradation. Surprisingly, this mechanism is shared with three recently reported compounds which all have unique chemical structures. ^42–44^ Thus, our discovery of the SW394703 MoA not only provides a unique scaffold with which to optimize a cyclin K degrader, but also provides a starting point for the design of neo-functional molecules that recruit alternative proteins to DDB1-containing cullin complexes for degradation. This idea draws inspiration from the IMiDs, a class of molecular glue degraders for which different chemical derivatives are able to selectively recruit diverse neo-substrates to the CRL4-CRBN E3 ubiquitin ligase. ^41^ Future studies focused on structural determination of the complex formed by SW394703, DDB1, and CDK12-cyclin K will be helpful in guiding the development of such neo-functional molecules. Combined, this study provides a framework for the generation and implementation of inducible mismatch-repair deficient cell lines as powerful tools for small molecule target investigation and biological discovery.

## Acknowledgements

We thank Anderson Frank for critically evaluating and editing our manuscript and helpful discussion, all current and former Nijhawan lab members for helpful discussions, Anastasia Beloglazkina for PVHD303 synthesis, members of the UT Southwestern high-throughput screening core. T.P.N. is supported by McKnight Fellowship. J.M.R. is supported in part by I-1612 (Welch) and RM1GM142002 (NIH). D.N. is a UT Presidential Scholar and holds the Joseph F. Sambrook, Ph.D. Distinguished Chair in Biomedical Science. J.K.D. holds the Julie and Louis Beecherl, Jr., Chair in Medical Science. J.M.R is the Ron Estabrook Distinguished University Chair in Biomedical Sciences. J.M.R., D.N., B.A.P., and D.G.M. acknowledge support of the NIH Moonshot program, U54CA231649. B.A.P acknowledges support from the UT Southwestern Simmons Comprehensive Cancer Center, 2P30CA142543-11. J.K.D. and D.N. and D.G.M acknowledge support of the Welch Foundation (grants I-1422, I-1879, and I-2040, respectively). The Program in Molecular Medicine is supported by an anonymous donor.

## Author contributions

T.P.N. performed confocal experiments, CDK12-DDB1 co-IP experiments, all MM201, MM202, and MM204 related experiments, and wrote the manuscript. M.F. performed all compound resistance screens. J.K. analyzed WES. Cell line engineering: iHCT116 (B.W., E.L. and T.P.N.); iS462 (T.P.N., C.C.); ATP6V1B2^D359N^ (V.K., N.B); PARP-1 KO and KI (M.G., G.R.); A673 ^TUBB4B T238A^ (J.M.P., D.G.M). V.K., N.B. and J.M.P. processed and analyzed barcode information. B.A.P. supervised high throughput screening of HCT116 cells. S.X. examined SW394703-dependent cyclin K degradation in HCT116. M.A. and J.M.R synthesized SW394703 and SW394704. Y.X. supervised WES analysis. J.K.D. and D.N. designed and supervised compound toxicity studies, target identification, target validation studies. D.N. conceived the study and wrote the manuscript.

---

**Supplement figure 1.**
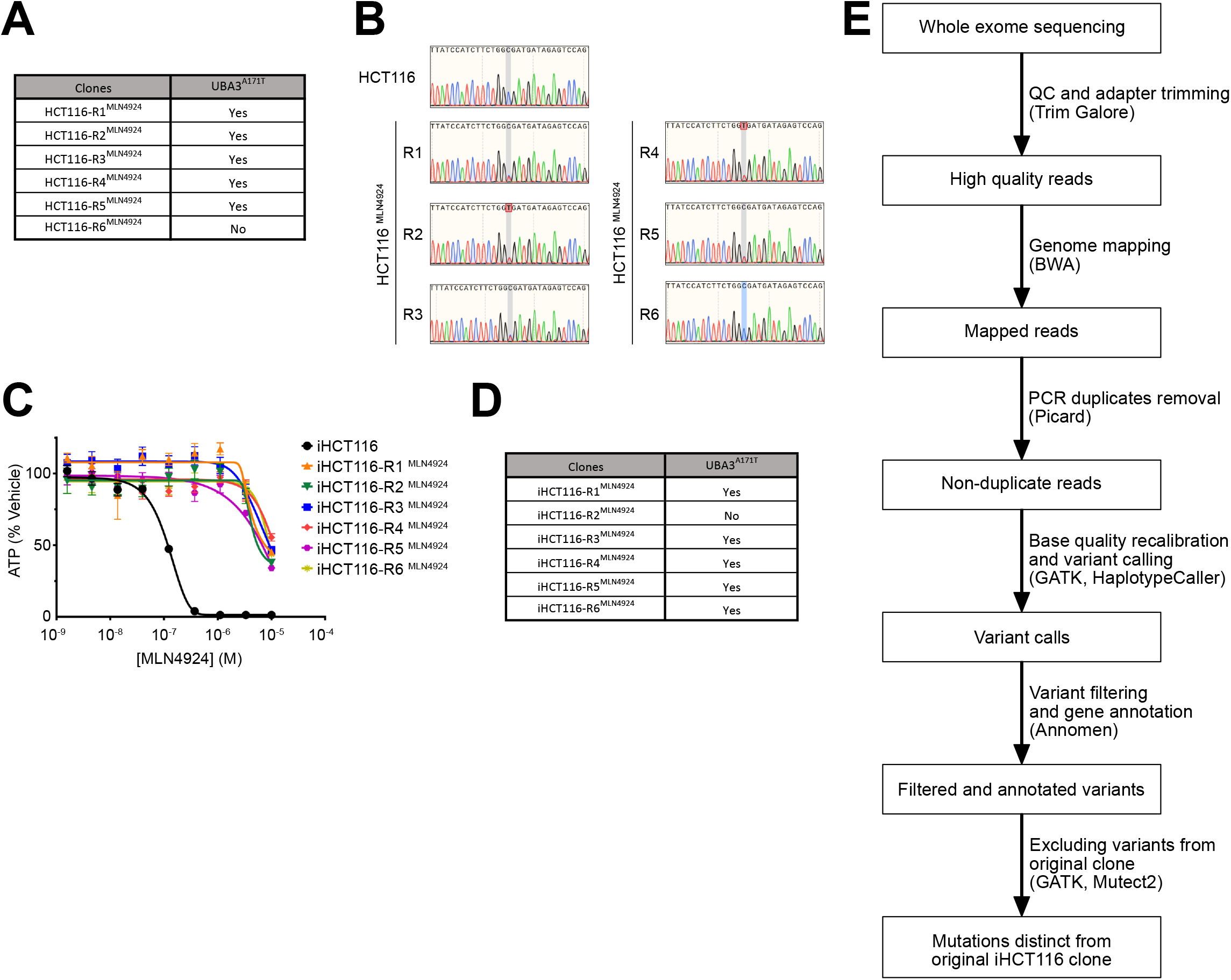
(A) Sanger sequencing results in 6 MLN4924 resistant clones raised from HCT116^WT^. (B) Sanger sequencing traces showing heterozygous UBA3^A171T^ mutation. (C) 72-hour growth assay of parental iHCT116 and 6 MLN4924 resistant clones raised from iHCT116 treated with MLN4924 in dose-response format. Points and error bars represent mean +/- SEM of three replicates. Error bars are not shown if SEM is smaller than the point. (D) Sanger sequencing results in 6 MLN4924 resistant clones raised from iHCT116. (E) Flowchart of whole-exome sequencing analysis.

**Supplement figure 2.**
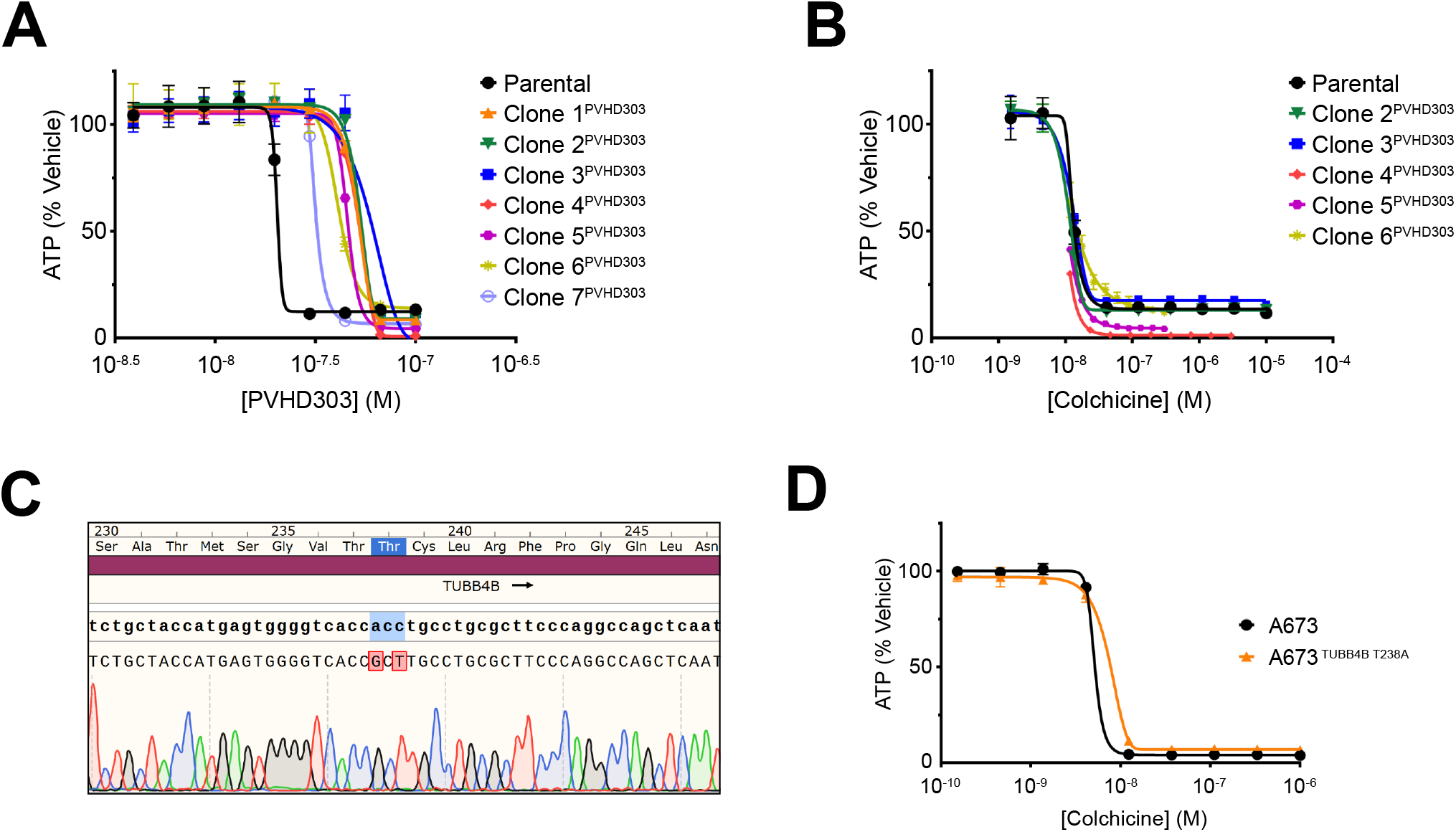
(A, B) 72-hour growth assay of parental iHCT116 and PVHD303 resistant clones treated with PVHD303 or colchicine in dose-response. (C) Sanger sequencing trace to show homozygous knock-in of TUBB4B^T238A^ mutation in A673. (D) 72-hour growth assay of A673 and A673 ^TUBB4B T238A^ treated with colchicine in dose-response. Points and error bars represent mean +/- SEM of three replicates. Error bars are not shown if SEM is smaller than the point.

**Supplement figure 3.**
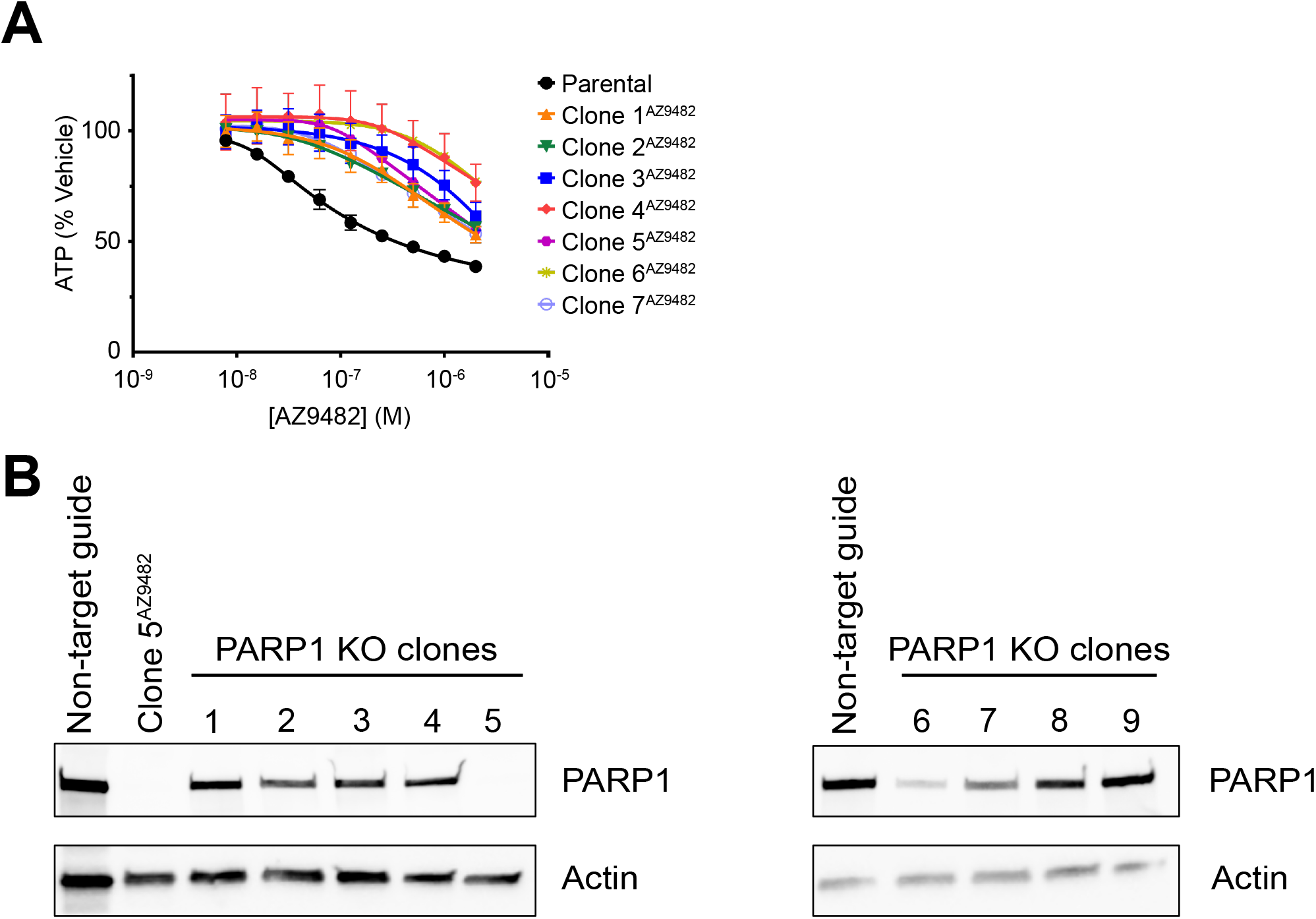
(A) 72-hour growth assay of parental iHCT116 and 7 AZ9482 resistant clones treated with AZ9482. (B) Western blot of lysates derived from iHCT116 infected with non-target guide or clones with guide targeting PARP1.

**Supplement figure 4.**
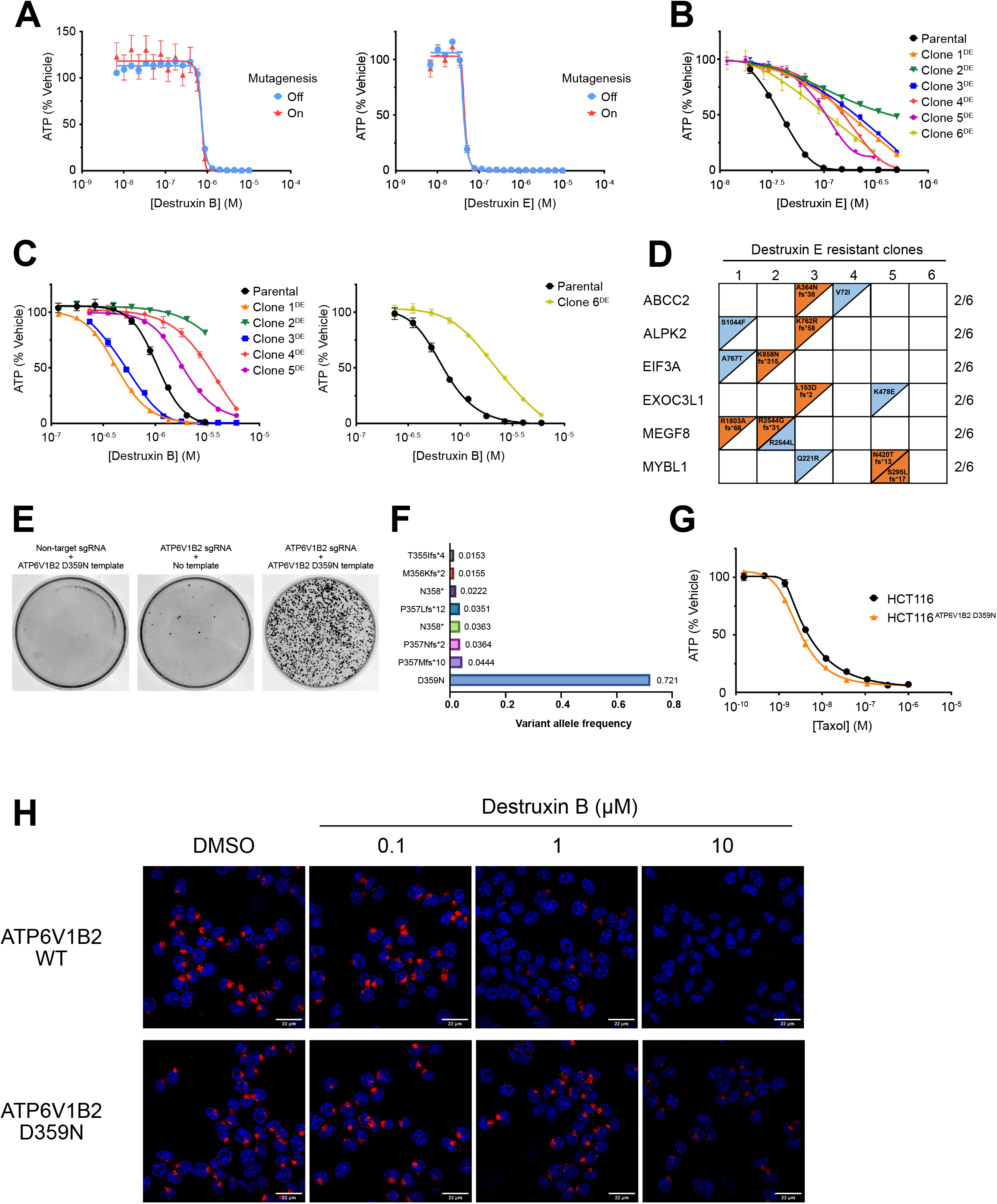
(A) 7-day growth assay of the Mut-off and on populations treated with DB (left), or DE (right) in dose-response. (B, C) 72-hour growth assay of parental iHCT116 and 6 DE resistant clones treated with DE (B) or DB (C). (D) WES analysis of DE resistant clones. (E) Crystal violet stain of transfected HCT116 treated with 97 nM DE for 2 weeks. (F) Amplicon sequencing result of the PCR product amplified from HCT116^ATP6V1B2 D359N^ population. (G) 72-hour growth assay of HCT116 and HCT116^ATP6V1B2 D359N^ treated with taxol in dose-response format. Points and error bars represent mean +/- SEM of two biological replicates. Error bars are not shown if SEM is smaller than the point. (H) Confocal images of HCT116 or HCT116^ATP6V1B2 D359N^ treated with bafilomycin A1 or destruxin B as indicated for 4 hours, followed by incubation of Hoechst reagent and lysotracker. Nuclei are stained blue. Lysotracker is red. Bar-scale represent 22 μm.

**Supplement figure 5.**
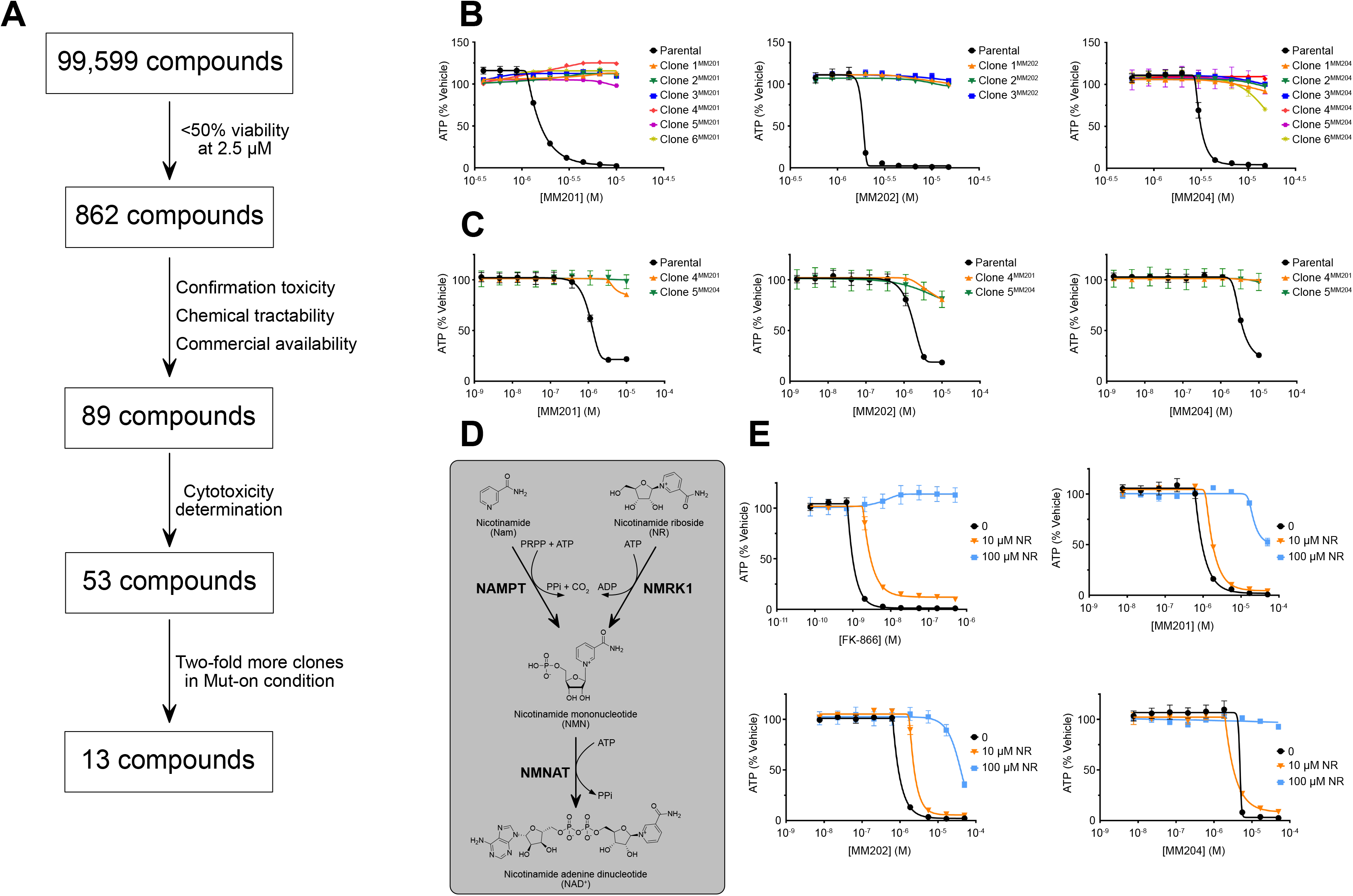
(A) High throughput chemical screen. (B) 72-hour growth assay of parental iHCT116 and MM201, MM202, and MM204 resistant clones treated with MM201, MM202, and MM204 in dose-response. (C) 72-hour growth assay of parental iHCT116, clone 4^MM201^, and clone 5^MM204^ treated with MM201, MM202, and MM204 in dose-response format. (D) Schematics of NAD^+^ generation pathways from nicotinamide (Nam) and nicotinamide riboside (NR) in cell. (E) 72-hour growth assay of parental iHCT116 treated with FK-866, MM201, MM202, and MM204 in dose-response, in the absence or presence of indicated concentration of NR. Points and error bars represent mean +/- SEM of two biological replicates. Error bars are not shown if SEM is smaller than the point.

**Supplement figure 6.**
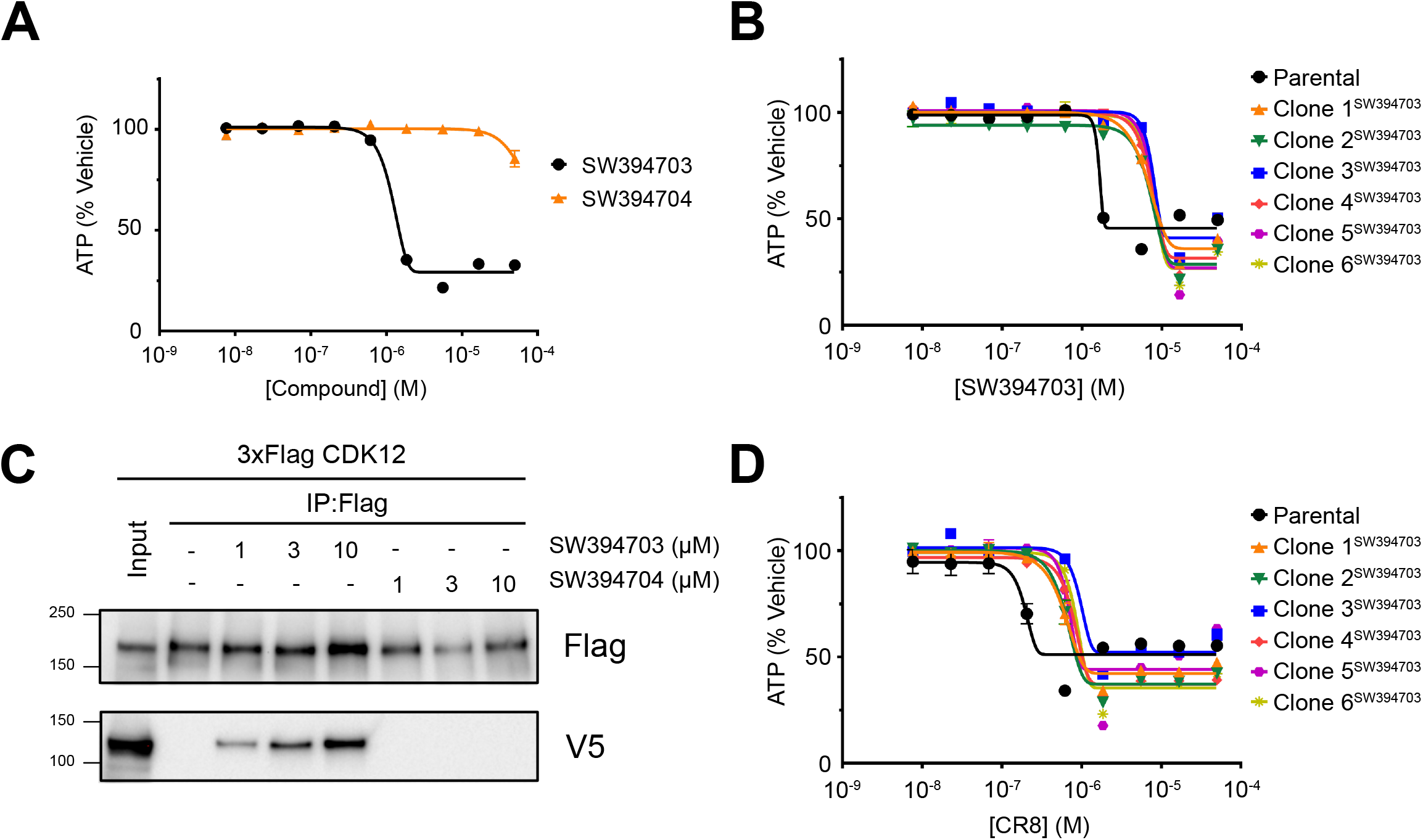
(A) 72-hour growth assay of HCT116 treated with SW394703 or SW394704 in dose response. (B, D) 72-hour growth assay of iHCT116 and SW394703 resistant clones treated with SW394703 (B) or CR8 (D). (C) Immunoprecipitation and western blot of 3xFlag expressing HCT116 lysates in the presence of SW394703 or SW394704.

**Supplement figure 7.**
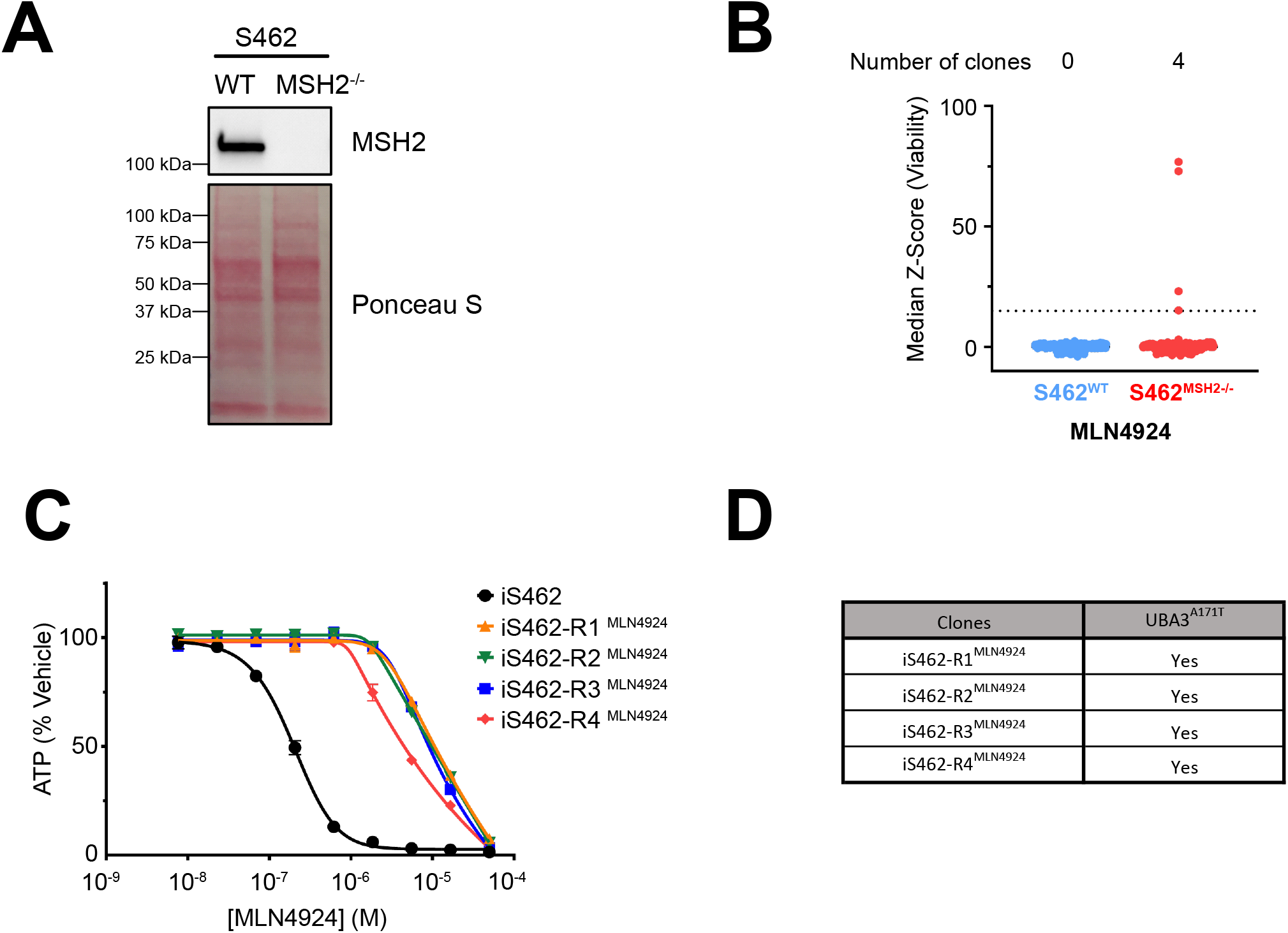
(A) SDS-PAGE and immunoblot analysis of wildtype S462 (S462^WT^) and MSH2 null S462 (S462^MSH2-/-^). (B) S462^WT^ and S462^MSH2-/-^ populations were subjected to 0.25 μM MLN-4924 selection for 2 weeks. Cells were plated in 96-WP, 3000 cells/well, 3 plates/population. At the end of the selection, cell viability in each well was assessed using alamarBlue. AlamarBlue signal was normalized and presented as median Z-score. Data points with Z-score value larger than 15 (above the dash line) were considered clones. (C) 72-hour growth assay of parental iS462 and 4 MLN4924 resistant clones raised from iS462 treated with MLN4924 in dose-response format. Points and error bars represent mean +/- SEM of two biological replicates. Error bars are not shown if SEM is smaller than the point. (D) Sanger sequencing results in 4 MLN4924 resistant clones raised from iS462.

## Materials and Methods

### Commercial chemicals

MLN-4924 (HY-70062) was purchased from MedChemExpress. AZ9482 (SYN-3046) was purchased from SYNKinase. MM201 (21512557), MM202 (81479831), and MM204 (17712729) were purchased from Chembridge. Nicotinamide riboside (SMB00907) was purchased from Millipore Sigma. Taxol (S1150) and FK-866 (S279902) were purchased from Selleck Chem. Colchicine (C9754) was purchased from Sigma-Aldrich. Destruxin B and Destruxin E were gifted by Hexagon Bio. Bafilomycin A1 (11038) was purchased from Cayman Chemical. CR8 (406333) was purchased from Medkoo bioscience.

### Chemical synthesis

PVHD303 was synthesized following the published procedure.

#### SW394703 and SW394704 synthesis

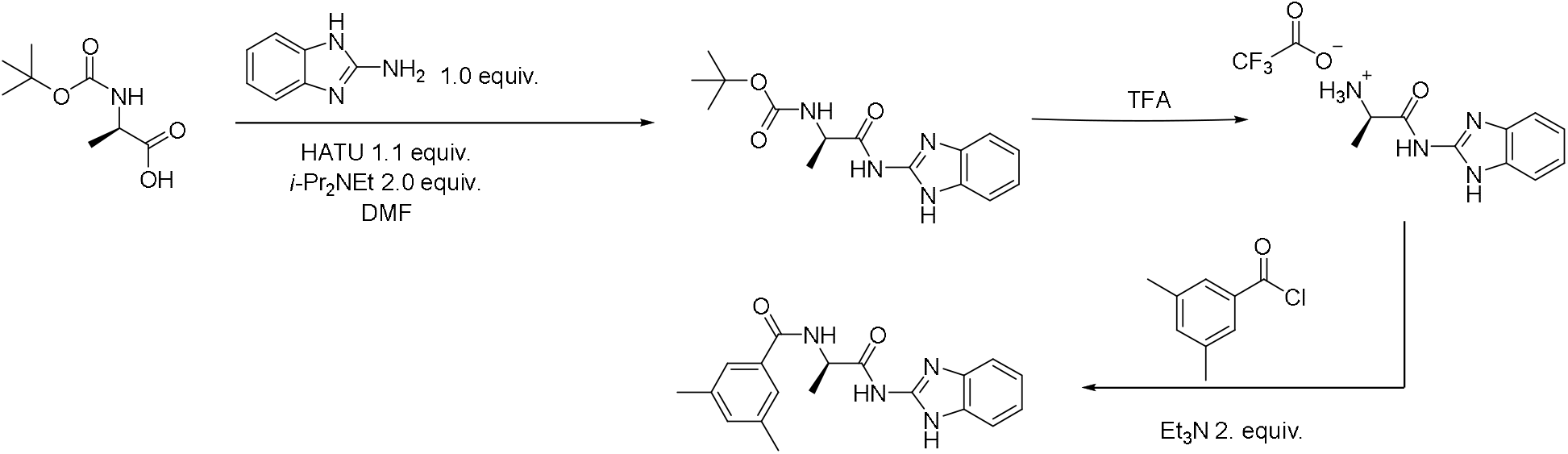

#### tert-Butyl (R)-(1-((1H-benzo[d]imidazol-2-yl)amino)-1-oxopropan-2-yl)carbamate

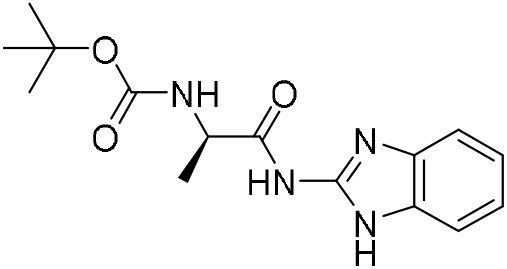

To a solution of (tert-butoxycarbonyl)-*D*-alanine (1 g, 5.3 mmol, 1.0 equiv) and 2-aminobenzimidazole (704 mg, 5.3 mmol, 1.0 equiv.) in DMF (20 mL) was added HATU (2.2 g, 5.3 mmol, 1.1 equiv) and DIPEA (1.5 mL, 10.6 mmol, 2.0 equiv). The reaction mixture was stirred at room temperature overnight. Once completed as judged by HPLC/MS, the desired product was obtained as a white solid in 71% isolated yield by filtration of the reaction mixture and used without additional purification. ^1^H NMR (400 MHz, CDCl_3_) δ 7.81 – 7.44 (m, 2H), 7.34 – 7.17 (m, 2H), 4.60 (s, 1H), 1.51 (d, *J* = 6.6 Hz, 3H), 1.46 (s, 9H). ESI-MS (m/z): 305.1 [M+H]^+^.

#### (R)-2-amino-N-(1H-benzo[d]imidazol-2-yl)propanamide TFA salt

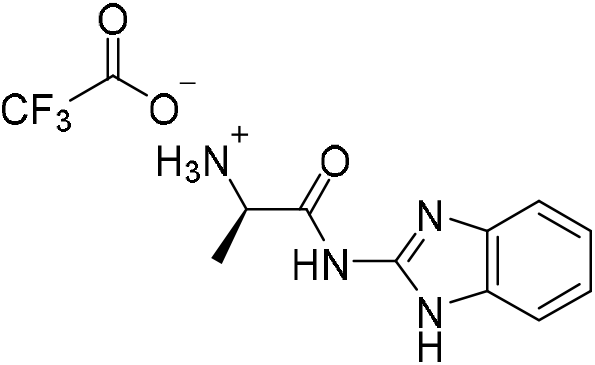

tert-Butyl (*R*)-(1-((1H-benzo[d] imidazol-2-yl)amino)-1-oxopropan-2-yl)carbamate (200 mg, 0.65 mmol) was dissolved in CH_2_Cl_2_ (2.5 mL) and 1.3 mL of trifluoroacetic acid was added at 0 °C. The reaction mixture was stirred at 0 °C for 1 h. Upon completion as judged by HPLC/MS, the reaction mixture was concentrated under a positive flow of nitrogen and used for the next step without further purification. ESI-MS (m/z): 205.1 [M+H]^+^.

#### SW394703 (*R*)-*N*-(1-((1H-benzo[d]imidazol-2-yl)amino)-1-oxopropan-2-yl)-3,5-dimethylbenzamide

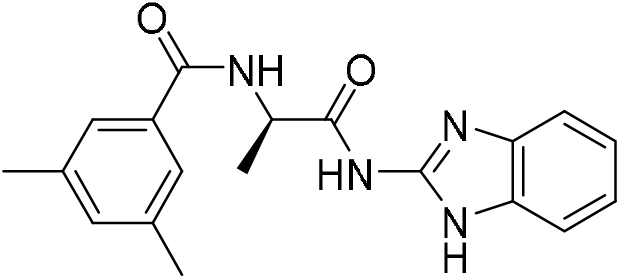

To a solution of 2-amino-N-(1H-benzo[d]imidazol-2-yl)propanamide TFA salt (crude product from the previous step) (50 mg, 0.16 mmol, 1.0 equiv) in 1.0 mL of CH_2_Cl_2_ was added Et_3_N (45 μL, 0.31mmol, 2.0 equiv) at 0 °C and the reaction mixture was stirred for 5 min at this same temperature 3,5-Dimethylbenzoyl chloride (25 μL, 0.17 mmol, 1.1 equiv.) was then added. After 30min of stirring at room temperature, the reaction mixture was concentrated and purified on silica gel (ISCO) to give product as a white solid in 63 % isolated yield. ^1^H NMR (600 MHz, CDCl_3_) δ 7.90 (s, 1H), 7.55 – 7.43 (m, 2H), 7.33 (s, 2H), 7.16 – 7.15 (m, 2H), 6.99 (s, 1H), 5.14 (p, *J* = 7.2 Hz, 1H), 2.12 (s, 6H), 1.55 (d, *J* = 7.2 Hz, 3H). ^13^C NMR (151 MHz, CDCl_3_) δ 175.27, 168.12, 147.48, 138.16, 133.42, 133.30, 125.07, 122.66, 50.45, 21.07, 18.46. ESI-MS (m/z): 337.1 [M+H]^+^.

#### SW394704

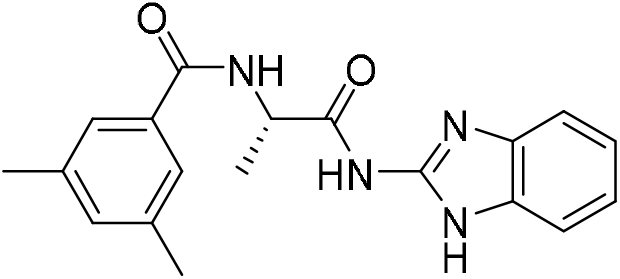

(*S*)-*N*-(1-((1H-benzo[d]imidazol-2-yl)amino)-1-oxopropan-2-yl)-3,5-dimethylbenzamide was prepared form (tert-butoxycarbonyl)-*L*-alanine following procedures described for compound **SW394703**.

### Antibodies

MLH1 (#11697-1-AP) antibody was purchased from Proteintech. PMS2 (sc-25315) and Actin (47778-HRP) antibodies were purchased from Santa Cruz Biotechnology. Flag-HRP (A8592) and V5-HRP (V2260) antibodies were purchased from Sigma. MSH2 (2017S), MSH6 (3996S), and PARP1 (#9542) antibodies were purchased from Cell Signaling Technologies. Cyclin K antibody (A301-939A-T) was purchased from Fortis Life Science.

### Cell culture, dose response curves, and viability assays

The identity of all human cell lines in this study was confirmed by short tandem repeat (STR) analysis. All cell lines were confirmed to be mycoplasma-free using a PCR based assay (Genatlantis).

Lenti-X HEK293T cells (Clontech, #632180), HCT116 (ATCC), and S462 cells are cultured in DMEM with high glucose (Sigma, #D5796), supplemented with 10% Fetal Bovine Serum (FBS) and 2 mM Glutamine. A673 (ATCC) are cultured in RPMI with high glucose (Sigma, #R8758), supplemented with 10% Fetal Bovine Serum (FBS) and 2 mM Glutamine. Cells were incubated at 37°C, 5% CO_2_.

For dose-response growth assay, 4000-5000 cells were plated in 96-well plate. The next day, cells were treated with compounds in dose response. All wells were normalized to 0.5% DMSO, and each dose was done in duplicates or triplicates. Cells were incubated at 37°C, 5% CO_2_, and 5% O_2_ for 72h. Cell viability was assayed using CellTiter-Glo^®^ (Promega, #G8462) per manufacturer instruction. Data were plotted, and IC_50_ values were calculated by baseline correction, curve fitting using a five-parameter dose-response curve in GraphPad Prism.

### SDS-PAGE and immunoblots

Cells were lysed in 1% SDS buffer A (50 mM HEPES pH 8.0, 10 mM KCl, 2 mM MgCl_2_) supplemented with 1 U/ml benzonase. Protein concentrations were measured by absorbance at 280nm and lysates were normalized. Samples were supplemented with 4x Laemmli sample buffer with 50 mM 2-mercaptoethanol before loaded on a Tris-Glycine gel. Gels were transferred onto 0.45 micron nitrocellulose membrane either by wet transfer or the Trans-Blot Turbo transfer system (Bio Rad). Membranes were stained with Ponceau S (0.1% w/v in 5% acetic acid) immediately after transfer for total protein visualization. Membranes were washed with Phosphate Buffered Saline with 0.025% Tween 20 (PBS-T) to remove Ponceau S. Membranes were then blocked with 5% non-fat dry milk diluted in PBS-T for 45 minutes. Blocked membranes were incubated with primary antibodies diluted in 5% bovine serum albumin in PBS-T overnight at 4°C. The next day, membranes were washed with PBS-T three times, 5 minutes each, before incubated with HRP-conjugated secondary antibodies diluted in 5% non-fat dry milk in PBS-T for 1 hour. Membranes were washed 3 times with PBS-T, 10 minutes each. Peroxidase signal visualized with ECL and imaged with Bio-Rad ChemiDoc.

### Plasmids

MLH1 cDNA was amplified from Hela cDNA library (5’ GAAGACACCGACTCTACTAGAGGATCTATTTCCGGTGAATTCATGTCGTTCGTGGCAGGG; 5’ GGGGGGAGGGAGAGGGGCGGGATCCGCGGCCGCTCTAGAACTAGTTTAACACCTCTCAAAGACTTTGTATAGATCAGG) and cloned into pLVX vector that was engineered to include Blasticidin resistant marker downstream of IRES element, tag-less or with 3xFlag and AID degron sequence at the C-terminus.

A plasmid containing *CDK12* isoform 2 (NM_015083.3) was obtained from UTSW Sequencing Core. CDK12 was PCR amplified using primers 5’- GGACGATGATGATAAAACTAGTCTAGAgcccaattcagagagacatgg and 5’- GAGGGAGAGGGGCGGGATCCggttagtaaggaactcctctccctct. The PCR amplicon was cloned into a pLVX-N-3xFlag vector which was derived from pLVX vector by Gibson Assembly. Plasmid was sequence verified.

### Lentiviral packaging and infection

Lenti-X HEK293T cells (Clontech, #632180) were transfected with 1 μg of the DNA construct of interest, 900 ng psPAX (Addgene, #12260), and 100 ng pMD2.G (Addgene, #12259) with TransIT-Lenti (Mirus, #MIR 6603) in a well of a 6-well plate, 5.5×10^5^ cells/well. Viral supernatants were collected 72h after transfection.

Infection in HCT116: Cells were infected with virus and 8 μg/mL polybrene (NEB, #H9268).

### Cell line generation

#### HCT116 overexpressing MLH1

Tagless MLH1 was packaged in lentivirus to infect HCT116. Infected cells were selected for blasticidin resistance.

#### iHCT116

HCT116 cells were infected with TIR1 and MLH1-3xFlag-AID viruses. Single clones were picked and assessed for AID-dependent degradation of MLH1. One clone was chosen and expanded for further experiments, named iHCT116.

#### iS462

Guide RNA targeting the genomic region surrounding MSH2 stop codon was cloned into pX330 (Addgene #42230). The repair template was constructed in a pGEM-T Easy vector to include 1 kilobase homology arms matching upstream and downstream sequences of the genomic locus. The repair template sequence contains 3xFlag and AID degron sequence, as well as an IRES Neo cassette flanked by two LoxP sites. S462 cells were transfected with the guide RNA and the repair template using TransIT LT (Mirus). Cells were selected with G418. Single clones were screen by western blot and PCR for homozygous MSH2-3xFlag-AID knock in. Knock-in clone was then infected with virus expressing TIR1.

### Generation of barcoded iHCT116

A random sequence of 24 nucleotides were conjugate to Neo-resistant gene by PCR (5’- TCTAGAGCGGCCGCGGATCCATGGTTATTGAACAAGATGGATTGCACGC, 5’- AGAGGTTGATTGTTCCAGACGCGTWSWSWSWSWSWSWSWSWSWSWSWSTCAGAAGAACT CGTCAAGAAGGCGATAGAAG), then cloned in pLVX vector. Assembled plasmids were transformed to electro-competent E. coli (Lucigen, #60242) and 3×10^6^ colonies resulting from the transformation were pooled together to generate a barcode plasmid DNA. The plasmid library was packaged into lentivirus to infect iHCT116 at a low multiplicity of infection (MOI) to achieve ~30% cell survival after neomycin selection. This resulted in a barcoded iHCT116 population in which most cells contain one unique barcode.

### Selection of resistant clones

Barcoded iHCT116 cells were cultured with or without 500 μM indole-3-acetic acid (IAA) for 2 weeks. Cells were plated in 96-well plates, 5000 cells/well. Compounds were added the next day by Tecan D300e digital dispenser. Medium containing compound of interest was replenished after 7 days. 14 days after compound selection, clones were screen by AlamarBlue viability assay. Clones were then expanded, and barcode genotyped to identify independent clones for further analysis.

### Sanger sequencing of UBA3

Genomic DNA isolated from MLN4924 resistant clones were used as template for amplification of a DNA sequence flanking alanine 171 region of UBA3 with primers 5’- ACGTGCAAATTGTTTTCTGCA and 5’- GGGCTTTAAAGGAGAAGGGG. PCR amplicons were Sanger sequenced.

### Barcode analysis

Genomic DNA was extracted from cell pellets using QiaAMP DNA Mini Kit (Qiagen, 51306). Barcodes were PCR amplified using 5’- GATATTGCTGAAGAGCTTG and 5’- CCAGAGGTTGATTGTTCCAGA. PCR products were sent for Sanger sequencing (individual) or Amplicon Sequencing (pool) at Genewiz.

### Next generation sequencing

Genomic DNA purification was performed using the QIAamp DNA Mini Kit (Qiagen). Whole exome sequencing was done at BGI. In short, genomic DNA libraries were prepared using the SureSelect Human All Exon V6 kit (Agilent Technologies). Exome libraries were subjected to next generation sequencing using DNBseq platform (MGI) with 150 bp paired-end reads.

### Whole exome sequencing analysis

Trim Galore (https://www.bioinformatics.babraham.ac.uk/projects/trim_galore/) was used for quality and adapter trimming. The human reference genome sequence and gene annotation data, hg38, were downloaded from Illumina iGenomes (https://support.illumina.com/sequencing/sequencing_software/igenome.html). The sequencing reads were aligned to the genome sequence using Burrows-Wheeler Aligner (BWA, v0.7.17).^57^ Picard (2.21.3) (https://broadinstitute.github.io/picard) was used to remove PCR duplicates and Genome Analysis Toolkit (GATK, 4.1.4.0) ^58,59^ was used to recalibrate base qualities. Calling variants and genotyping were performed using GATK HaplotypeCaller and the variant calls were filtered by applying the following criteria: QD (Variant Confidence/Quality by Depth) < 2, FS (Phred-scaled p-value using Fisher’s exact test to detect strand bias) > 60, MQ (RMS Mapping Quality) < 40, DP (Approximate read depth) < 3, GQ (Genotype Quality) < 7. Custom Perl scripts (https://github.com/jiwoongbio/Annomen) were used to annotate variants with human transcripts, proteins, and variations (RefSeq and dbSNP build 151), calculate variant allele frequencies. To date, we have sequenced 158 independent clones resulting from screening several distinct compounds and found 104 genes harboring any mutation (including synonymous, non-synonymous, non-coding, nonsense, and insert-deletion) in 25% of all clones (Supplemental Table 1). Some of these genes harbored repetitive sequences that may have contributed to a high mutation rate. We considered these mutations to be a consequence of this artifact rather than related to compound resistance, and removed these genes from further analysis.

### Knock-in of TUBB4B T238A allele in A673 cells

We performed homology-directed repair using Alt-R CRISPR-Cas9 System and ultramer oligo from Integrated DNA Technologies (IDT) in A673 Ewing sarcoma cells to knockin T238A mutation in TUBB4B. To prepare the gRNA complex, we combined 10 μL of Alt-R CRISPR-Cas9 crRNA (100 μM) (sequence: 5’- CUG GCC UGG GAA GCG CAG GCG UUU UAG AGC UAU GCU-3’) and 10 μL Alt-R CRISPR-Cas9 tracrRNA (100 μM). The mixture was heated to 95°C for 5 minutes and then slowly cooled to room temperature. The ribonucleoprotein complex with Cas9 was formed by combining 9 μL of gRNA complex with 6 μL Alt-R Cas9 enzyme and incubating at RT for 10-20 minutes. Two million A673 cells were resuspended in 120 μL of SF Cell Line 4D-NucleofectorTM X Kit L (Cat. #: V4XC-3024). The transfection mix was prepared using: 15 μL of RNP complex, 3.6 μL of 100 μM Ultramer ssODN donor (sequence: 5’- C*C*T GAA CCA CCT GGT GTC TGC TAC CAT GAG TGG GGT CAC CGC TTG CCT GCG CTT CCC AGG CCA GCT CAA TGC TGA CCT GCG GAA G*C*T-3’; *signify phosphorothioate modified bases) and added to 60 μL of previously prepared cell suspension. Nucleofection was performed using 4D-Nucleofector™ core unit from LONZA. Cells were plated into one well of a 6-w plate after nucleofection in 2 mL of RPMI (5% FBS). After 48 hours, sham cells or ssODN (TUBB4BT238A) cells were plated into a 6-w plate. Each sample was plated into a well of a 6-well plate using 0.5 million cells per well. Cells were treated with compound 21 from Povedano et al. [PMID 33180487] at 400 nM. Media was replenished every 3 – 4 days over the course of 2 weeks followed by expansion in media without compound for 1 week. After compound 21 selection, individual resistant clones were sequenced to identify the A673-TUBB4BT238A/T238A clone used in this manuscript.

### Knock-in of PARP1 Y896C and A898T allele

Guide RNA targeting PARP1 (5’- AAACATGTAGCCTGTCTGGA) was cloned into pX330 vector (Addgene, #42230). HCT116 cells were transfected with the guide RNA and single-stranded oligo 5’- tcagtgaacagctgctcctaatttcccatcagaagattctcagctctcccttttccgaccttccagacaggctacatgtttggtaaagggatcta tttcactgatatggtctccaagagtgccaactactgccatacgtctcagggagacccaataggcttaatcctgttgggagaagttgcccttggaaacatgtg agt for A898T mutation and 5’- tggcactcagtgaacagctgctcctaatttcccatcagaagattctcagctctcccttttccgaccttccag acaggctacatgtttggtaaaggaatctgtttcgctgacatggtctccaagagtgccaactactgccatacgtctcagggagacccaataggcttaatc ctgttgggagaagttgcccttggaaacat for Y896C mutation using TransIT-LT (Mirus). Afterward, the populations were treated with 350 nM AZ9482 for 2 weeks to select for cells with PARP1 A898T and Y896C knock-in mutations.

### Knock-out of PARP1

Guide RNA targeting PARP1 (5’- CCACCTCAACGTCAGGGTGC) was cloned into LentiCRISPRv2 vector (Addgene, 52961). Lentivirus carrying CRISPR/Cas9-PARP1 guide was transduced to HCT116 in the presence of 8 μg/ml polybrene. Cells were treated with 2 μg/ml puromycin to select for infected cells. Infected population was later subcloned. All clones were screened for PARP1 protein expression by western blot.

### Knock-in of ATP6V1B2 D359N allele

Guide RNA targeting ATP6V1B2 (5’- AAACTTACCATCATTAGGCA) was cloned into pX330 vector (Addgene, #42230). HCT116 cells were transfected with the guide RNA and single-stranded oligo 5’- tacatgtatacagatttagccacgatatatgaacgcgctgggcgagtggaagggagaaacggctcgattactcaaatccctattctGacAatgcctaatA atggtaagttttggtatttggattataacacacctaatcattttaaagagagagaagacccatctaccttttcgagttgaagttattttgagaacacatc using TransIT-LT (Mirus). Afterward, the cell population was treated with 96 nM Destruxin E for 2 weeks to select for cells with ATP6V1B2 D359N knock-in. All clones were pooled together to make HCT116^ATP6V1B2 D359N^ population for further experiments.

### Confocal experiments

HCT116 and HCT116^ATP6V1B2 D359N^ cells were plated on the 8-chambered coverslip (Ibidi, #80827) the day before treatment. Cells were treated with 0.2 μM bafilomycin A1 and various concentrations of Destruxin B and E for 4 hours. Afterward, cells were incubated with 10 nM Lysotracker (Thermo Fisher, L12492) and 1 μg/ml Hoechst (ThermoFisher, H3570) for 45 minutes. Cells were imaged using Zeiss LSM880 Airyscan.

### High throughput chemical screen

HCT116 cells were plated in 384-well plates at 700 cells/well in a total volume of 60 μl. The next day, compounds were administered at 2.5 μM via BiomekFX. 96 hours after compound addition, cell viability was assayed using CellTiter-Glo. A potential hit was picked if its corrected value was less than −3 standard deviation from the mean of all compounds in a run, and the viability was less than 50% compared to DMSO treatment. In the confirmation screen, 862 hits from the primary screen were re-tested at 1, 3, 10 μM in a 96-hour assay. Compounds with viability less than 50% at 10 μM treatment were further subjected to cherry-picking based on their chemical tractability and commercial availability. 89 compounds were resupplied and tested in a 7-day viability assay. 53 cytotoxic compounds were applied to iHCT116 system to look for compounds that yielded mutagenesis-driven resistant clones.

### Co-immunoprecipitation of 3xFLAG-CDK12 and DDB1-V5

Anti-FLAG M2 antibody (Sigma-Aldrich, F3165) was coupled to magnetic beads (Dynabeads M-270 Epoxy, NEB 14302D) at the ratio of 10 μg antibody/mg of beads in the presence of 1 M ammonium acetate and 0.1 M sodium phosphate, pH 7.4 at 37°C overnight. HCT116 cells stably expressing EV or 3xFLAG-CDK12 was transfected with DDB1-V5 for 24-48 hours prior to the experiment. Cells were lysed in lysis buffer (50 mM HEPES, pH 7.4, 300 mM NaCl, 0.1% Triton-X100). The resulting lysates were centrifuged at 15,000 g for 10 min at 4°C. Clarified lysates were incubated with SW394703, SW394704 or DMSO as indicated for 30 min at 4°C. Afterward, 0.2 mg of anti-FLAG-conjugated magnetic beads was mixed with the clear lysates for 15 min on a rotating platform at 4°C, followed by 3 washes with binding buffer (50 mM HEPES pH 7.4, 300 mM NaCl, 0.5% NP-40). Bound proteins were eluted with 1 mg/ml 3xFLAG peptide.

