## Supplemental Table 1 for "Inducible mismatch repair streamlines forward genetic approaches to target identification of cytotoxic small molecules"

| | Gene | Number of clones with mutations out of 158 sequenced clones |
|---|---|---|
| 1 | ANKRD36C | 158 |
| 2 | KIR2DL4 | 158 |
| 3 | MUC3A | 158 |
| 4 | MUC6 | 158 |
| 5 | TAS2R19 | 158 |
| 6 | MUC19 | 158 |
| 7 | OR8U1 | 158 |
| 8 | LMLN2 | 158 |
| 9 | KRTAP5-7 | 158 |
| 10 | MUC16 | 155 |
| 11 | KIR3DL2 | 154 |
| 12 | GOLGA6L2 | 154 |
| 13 | TRIOBP | 154 |
| 14 | HLA-DRB1 | 153 |
| 15 | LILRB1 | 153 |
| 16 | KRTAP1-3 | 152 |
| 17 | KRT18 | 148 |
| 18 | TPSD1 | 145 |
| 19 | ZNF717 | 145 |
| 20 | LOC101928804 | 144 |
| 21 | ANKRD36 | 143 |
| 22 | MIR7155 | 143 |
| 23 | TAS2R46 | 141 |
| 24 | NPIPB15 | 139 |
| 25 | MUC17 | 135 |
| 26 | MLH1 | 131 |
| 27 | HLA-DQA2 | 130 |
| 28 | FADS6 | 129 |
| 29 | NBPF20 | 123 |
| 30 | FAM166B | 123 |
| 31 | HRNR | 111 |
| 32 | MIR558 | 109 |
| 33 | PCDHA8 | 105 |
| 34 | SON | 105 |
| 35 | SLC25A5 | 104 |
| 36 | PER3 | 103 |
| 37 | GOLGA8B | 102 |
| 38 | IGFN1 | 101 |
| 39 | MAP3K4 | 99 |
| 40 | UBR2 | 93 |
| 41 | PRB3 | 90 |
| 42 | RBFOX1 | 88 |

| | | |
|---|---|---|
| 43 | SLC9B1 | 88 |
| 44 | PPIP5K2 | 87 |
| 45 | CNOT1 | 87 |
| 46 | ACACA | 83 |
| 47 | ANKRD30B | 82 |
| 48 | CEP170B | 82 |
| 49 | DGKE | 82 |
| 50 | ANKRD30A | 81 |
| 51 | PCDH12 | 80 |
| 52 | ZNF880 | 77 |
| 53 | KAT2A | 76 |
| 54 | KRTAP9-9 | 74 |
| 55 | THAP12 | 73 |
| 56 | HTT | 73 |
| 57 | NUTM2E | 71 |
| 58 | ANKLE1 | 69 |
| 59 | MAGEC1 | 68 |
| 60 | HYDIN | 65 |
| 61 | GXYLT1 | 64 |
| 62 | ERN2 | 62 |
| 63 | SPRED3 | 62 |
| 64 | CNN2 | 61 |
| 65 | MUC21 | 61 |
| 66 | PIN4P1 | 61 |
| 67 | KRTAP4-7 | 60 |
| 68 | ZNF626 | 60 |
| 69 | JPH3 | 59 |
| 70 | MUC20 | 58 |
| 71 | TNRC18 | 57 |
| 72 | TTC21B | 57 |
| 73 | RPL13AP6 | 56 |
| 74 | TTN | 55 |
| 75 | EVI5 | 55 |
| 76 | LILRB3 | 54 |
| 77 | OR2T4 | 52 |
| 78 | KRTAP5-5 | 52 |
| 79 | CCDC40 | 51 |
| 80 | MEOX2 | 50 |
| 81 | OR4N3P | 50 |
| 82 | THAP11 | 50 |
| 83 | NACAD | 49 |
| 84 | MAML2 | 49 |
| 85 | PCMTD2 | 48 |

| | | |
|---|---|---|
| 86 | HLA-B | 47 |
| 87 | HRC | 47 |
| 88 | KCNN3 | 46 |
| 89 | GPR21 | 45 |
| 90 | PAX6 | 45 |
| 91 | ZIC5 | 45 |
| 92 | ATXN7L1 | 44 |
| 93 | PHGR1 | 44 |
| 94 | NAP1L2 | 43 |
| 95 | SLC5A4 | 43 |
| 96 | KRT10 | 42 |
| 97 | CDC14C | 41 |
| 98 | DNHD1 | 41 |
| 99 | FOXN4 | 41 |
| 100 | PARG | 40 |
| 101 | FBRSL1 | 40 |
| 102 | ZFTA | 40 |
| 103 | FRMPD3 | 40 |
| 104 | PAXIP1 | 40 |
